# Computational exploration of molecular receptive fields in the olfactory bulb reveals a glomerulus-centric chemical map

**DOI:** 10.1101/489666

**Authors:** Jan Soelter, Jan Schumacher, Hartwig Spors, Michael Schmuker

## Abstract

Progress in olfactory research is currently hampered by incomplete knowledge about chemical receptive ranges of primary receptors. Moreover, the chemical logic underlying the arrangement of computational units in the olfactory bulb has still not been resolved. We undertook a large-scale approach at characterising molecular receptive ranges (MRRs) of glomeruli innervated by the MOR18-2 olfactory receptor in the dorsal olfactory bulb (dOB). Guided by an iterative approach that combined biological screening and machine learning, we selected 214 odorants to characterise the response of MOR18-2 and its neighbouring glomeruli. We discovered several previously unknown odorants activating MOR18-2 glomeruli, and we obtained detailed MRRs of MOR18-2 glomeruli and their neighbours. Physico-chemical MRR descriptions revealed that the spatial layout of glomeruli followed a chemical logic. Our results confirm earlier findings that demonstrate a partial chemical map underlying glomerular arrangement in the dOB. Moreover, our novel methodology that combines machine learning and physiological measurements lights the way towards future high-throughput studies to deorphanise and characterise structure-activity relationships in olfaction.

## Introduction

For most sensory systems, the receptive fields of primary receptor neurons are fairly well identified, for example the spectral tuning of photoreceptors, or frequency tuning of cochlear hair cells. Moreover, topographical arrangements like retinotopy in vision, tonotopy in audition, and somatotopy in somatosensation have been described ^1–3^, which minimize wiring length and likely improve the efficiency of computation ^4–6^.

In the olfactory system, the organisation of the main olfactory epithelium (MOE) in zones that define preferred locations of olfactory sensory neuron (OSN) is an instance of such topography ^7–9^. Some zonal co-localisation of OSNs with similar molecular receptive ranges (MRRs) has been observed ^10–12^, but well-defined zonal adherence could only be observed for a subset of receptors. It has been suggested that zonal OSNs expression patterns propagate to the olfactory bulb (OB) ^13^.

Conflicting interpretations of experimental results exist however as to whether glomerular positions in the OB reflect a *chemical* topology, in the sense that spatial proximity of glomeruli indicates proximity of their preferred ligands in chemical space. Mori et al. ^14^ collected information from a large body of literature about tuning and spatial arrangement of glomeruli in the mouse and found that glomeruli were apparently arranged in a patchy map of (quote) “molecular-feature clusters are located at stereotypical positions in the OB”, and that “the glomerular sheet represents the characteristic molecular features in a systematic gradual, and multidimensional fashion”. Similarly, Johnson and Leon reported that odorants sharing structural chemical features such as functional groups, aromatic rings and solubility evoked clustered responses ^15^, that shift in a systematic way according to carbon chain length, confirming earlier observations ^16^.

Contrarily, other reports dispute that chemical similarity is reflected in the bulbar location of glomeruli. Soucy et al. found that neighbouring glomeruli tend to respond to overlapping sets of ligands, but dismiss the idea of a chemotopic ordering, claiming that the intrinsic variability of glomerulus placement is greater than the accuracy of the chemical map ^17^. They also shine a critical light on Mori’s hypothesis on chemotopic modules as they observe glomeruli in a region apparently tuned to aldehydes that do not respond to that substance class at all. Generally, they could not confirm any chemotopic arrangement beyond a few glomerular radii.

Ma and colleagues ^18^ reported that no chemotopic organisation could be observed in a large dataset of glomerular responses in the dOB using about 60 odorants. However, their analysis mainly rests on functional group comparisons and largely ignores other molecular features, therefore their assessment does not reflect all aspects of chemical similarity. Their report was opposed by Yablonka and coworkers ^19^, who pointed out flaws in their methodology regarding the normalisation of descriptors, and proposed an analysis that indeed suggested a relation between spatial distance of glomeruli and chemical distance of their preferred ligands.

These seemingly conflicting reports reveal the general problem in assessing chemotopy, namely that no generally valid, rigorous definition of chemical similarity exists. For example, similarity of odorants can be described by physico-chemical properties ^20,21^. Likewise, molecular similarity can be assessed by vibrational spectra, which have been used to successfully predict receptor affinity, as originally proposed in drug discovery ^22^, and more recently confirmed in insect olfaction ^23^.

An important distinction to make when considering the seemingly contradictory evidence is whether actual chemoptopy has been assessed by calculating the overlap of MRRs in chemical space, or rather *tunotopy*, that is, merely computing the amount of overlap in the ligand spectra ^18^. Tunotopy, while straightforward to compute empirically, lacks an immediate connection to physico-chemical space, thus preventing further insights into the chemical map on the OB surface. This chemical map requires an abstraction away from discrete compounds, and towards physico-chemical properties that define the map “coordinates”. Due to the high dimensionality of chemical space, a reliable assessment of chemotopy thus critically depends on the number of odorants and glomerular responses taken into account.

MRR data are still sparse, and the sheer number of OR genes that are still to be de-orphaned and characterised has so far impaired a comprehensive view on chemotopy in the OB. Chemical tuning of a glomerulus and its neighbourhood is rarely assessed using more than a hundred odorants, which provides a tiny window into the vast and extremely high-dimensional olfactory chemical space. Detailed MRRs are only available for a small amount of receptors ^24–27^, partial MRRs merely for a few more ^20,28–33^. The scarcity of data limits the explanatory power that can be expected from any approach that pursues the investigation of chemotopy organisation in the OB that relies on comparisons of a large number of glomerular MRRs.

In this study we address the scarcity of data by automated biological in-vivo screening to deorphanize glomeruli, characterise their MRRs, and we leverage machine learning to assess tunotopic and chemotopic relations between neighboring glomeruli. We started by mapping the MRR of a single glomerulus as completely as possible, and subsequently explored the chemotopic embedding in its neighbourhood. This contrasts earlier attempts that aimed to investigate the topographic relationship of many glomeruli with fragmentary MRRs.

We initially focused on a glomerulus in the dOB, the class-1 receptor MOR18-2 (a.k.a. MOL2.3, or OLFR78), that was easily accessible by imaging and for which a genetic marker for GFP labelling the corresponding olfactory sensory neurons (OSNs) existed. We then explored MRRs of adjacent glomeruli, finally extending the analysis on the whole area covered by imaging, assessing chemotopy on several spatial scales. We used machine learning to obtain a physico-chemical activation model of MOR18-2, thus defining its MRR by means of physico-chemical properties. This allowed us to map the location of a glomerulus in chemical space to its location on the dOB, getting further insight into chemotopy.

## Methods

Mice were anaesthetized using urethane (1.5 g/kg i.p.). Anaesthetic was supplemented throughout the experiments and the body temperature was kept between 36.5C and 37.5C using a heating pad and a rectal probe. For imaging a craniotomy over one or both olfactory bulb was cut. The *dura mater* was removed and the imaging chamber was filled with agar (1.5 %) and covered with a glass cover slip. The prepared skull was fixated with cement to a metal plate under the microscope. All animal care and experimental procedures were approved by the regional authorities and carried out in accordance with the animal ethics guidelines of the Max Planck Society, approved by the regional council (“Regierungspräsidium”) Darmstadt, Germany.

Odorants were filled into glass vials under Argon atmosphere in order to avoid any contamination. A two-arm robot (Combipal, CTC-Analytics, Zwingen, Switzerland) using the Software Chronos (Axel Semrau, Sprockhoevel, Germany) was used to sample 2.5 ml of the odour headspace with a pre-heated (45ºC) syringe and inject it (intrinsic optical signal experiments: 0.5 ml/s; SpH experiments: 1 ml/s) into a constant carrier flow of filtered and humidified air (2 l/min) into the mouse’s nose. Odour molecules reached the nose 2.5 s ± 0.3 s after recording onset as measured by a photoionization detector (Aurora Scientific, Canada). We measured the response to different stimuli sets of 4 to 53 odours in 78 MOL2.3-IGITL mice ^34^, 3 heterozygote OMP-SpH mice ^35^ and 15 heterozygote MOL2.3-IGITL OMP-Sph cross-breeds. Each stimulus set was at least measured twice in each mouse with odours presented in a pseudo-randomized sequence. We used mice of both sexes, aged 5-23 weeks.

After each odour presentation the syringe used for odour transfer was flushed with nitrogen for 72 s to minimize contamination. The sequence of odour presentation was pseudo-randomised to further quell any systematic influence of potential contamination. In addition, we recorded periods when no odorant was presented at the mean point between two measurements. Those ‘blank’ periods were later used for bleaching correction in SpH-imaging (see below).

### Intrinsic Optical Signal (IOS) Imaging

Odour responses were recorded in the dorsal olfactory bulb for 12s at 5Hz using a macroscope (Pentax zoom lens 12-48 mm, f = 1:1.0 and Nikkor lens 135 mm, f = 1:2.0) and an Orca-R2 camera (Hamamatsu, Japan; 1024 × 1344 px) under illumination with red light (690 nm) with a frame rate of 5 Hz (60 frames in 12 seconds). Uni-lateral OB recordings were performed at a focal length of 24 mm (field of view 1.63 mm × 1.24 mm), whereas bilateral recordings were performed at a focal length of 48 mm (field of view 3.26 mm × 2.48 mm). Before and after each presentation of the entire stimulus set, the pattern of blood vessels was recorded using green illumination (546 nm, ‘green image’) and controlled for shifts to exclude movement artefacts. Furthermore, the MOR18-2 glomeruli were located using blue illumination (475 nm, ‘GPF image’) and an emission filter at 535 nm.

To increase signal-to-noise ratio and reduce computational load we binned the raw data with an 8 × 8 px spatial and a 12 frame temporal window. Then the odour induced activation was calculated as the relative decrease of reflectance ∆*R*/*R* = −(*R* − *R*_0_)/*R*. *R*_0_ was the mean reflectance during the first 2 s after recording onset, before the odours reached the nose (which happened at 2.5 s ± 0.3 s, see above). Furthermore the data was spatially bandpass-filtered with two Gaussian filters (σ_*low*_ = 10 px, σ_*high*_ = 1 px) and spatially down-sampled by a factor of 2. The final resolution of the measurement time series was thus 64 × 84 px (with each pixel being 19.4 × 19.4 µm) at 0.42 Hz. The concatenation of the pre-processed frames for all odours lead to the measurement matrix **Y** ∈ *R*^*F×P*^ with element *Y*_*f,p*_ being the observed value of the *p*^*th*^ pixel in the *f*^*th*^ frame.

### Synapto-pHluorin (SpH) Imaging

Synapto-pHluorin imaging was performed with a 2-photon laser scanning microscope (Prairie Technologies, Middleton, TN, USA), a 16x water immersion objective (N.A. 0.8, Nikon, back aperture overfilled) and a MaiTai DeepSee laser (50-170mW, tuned to 880nm, 80 MHz repetition rate of pulses 120fs in length; Spectra-Physics/Newport, Santa Clara, CA, USA). Emitted light was separated and recorded into a green 525 nm and red 607 nm channel.

Functional imaging (128 × 128 pixel, 5.02 µm/pixel) was performed for each odour 12.77 s with 94 Hz at a fixed z-position. To increase signal-to-noise ratio and reduce computational load we binned the raw data in time to 120 frames, corresponding to a frame rate of 0.94 Hz. Then the odour induced activation was calculated as the increase in fluorescence *F* − *F*_0_. *F*_0_ was the mean fluorescence on the first 2 s after recording onset. Furthermore, the data was low pass filtered (Gaussian filter σ = 1.5 px) and down-sampled by a factor of 2. The final resolution of the measurement matrix **Y** was thus 64 × 64 px at 0.94 Hz. To account for bleaching effects, the same procedure was applied to control measurements without any odour stimulation. Out of these we calculated for each pixel the mean bleaching time course and subtracted those from the pixels’ odour responses time courses.

Before and after functional imaging we acquired anatomical images (512 × 512 pixel, 1.25 µm/px) at 3µm steps in z-direction. After functional imaging the mice were first killed by an overdose of urethane. Subsequently the OB was cleaned with artificial cerebro-spinal fluid (aCSF), fixated with methanol (1 min), rinsed for 10 min with buffer A (PBS 100 mM, MgCl2 2mM, EGTA 5 mM), 10 min with buffer B (PBS 100mM, MgCl2 2mM, Na-deoxycholate 0.01%, Igepal 0.02%) and finally stained by Fast Red Violet (FRV) before anatomical images were obtained. This made MOR18-2 glomeruli visible in the red channel (607 nm, ‘FRV image’).

### Glomerular response spectra

We segmented the functional image series into individual glomeruli and their activation course by means of regularized non-negative matrix factorization (rNMF) ^36^. Such a factorization disaggregates the measurement matrix **Y** into *k* components with a spatial signal distribution **x**_*k*_ and a common activation course **a**_*k*_ of the participating pixel. The factorization contained spatial smoothness (governed by parameter α_*sm*_) and sparseness regularization (α_*sp*_) to promote a disaggregation into spatial distinct but possibly overlapping components, i.e. glomeruli arranged side by side with an overlapping signal distribution.

We decomposed IOS image series into 150 components with fixed smoothness regularization parameter α_*sm*_ = 2. The sparseness regularization parameter α_*sp*_ was adjusted such that spatial component correlation was just below 0.5 (see Soelter et al., 2014, for a justification of this value). In image analysis performed to extract MOR18-2 responses, we dismissed the usual non-negativity constraint on the components’ activation courses to capture possible odour induced inhibitions. In contrast, in analysis to extract response spectra across the full dorsal OB we kept the non-negativity constraint as it increased the trial-to-trial correlation of response spectra.

SpH image series were disaggregated by the same approach. With respect to the reduced field of view, the factorization was performed only into 20 components with an extended smoothness regularization of α_*sm*_ = 5.

Odour responses 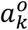 of each component *k* (e.g., glomeruli) were obtained by averaging the activation of frames 3 and 4 in IOS (4.8-9.6 s after recording onset) and frames 8-11 (6.6 - 10.3 s) in SpH measurements. All odour responses of a measurement composed the component’s response spectrum 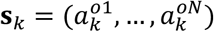. To remove non-glomerular components of the factorization, we only took components into account that exhibited a trial-to-trial correlation above 0.6 between response spectra of odour set repetitions within an animal ^36^. Components were manually assigned as MOR18-2 glomeruli if their pixel participation matched the GFP marked location of MOR18-2. The overall response strength varied strongly across animals, probably due to individual differences in glomerular layout – some animals only had one dorsal instance or MOR18-2, others up to four. Therefore, we normed MOR18-2 responses to its Methyl propionate (MP) response in the same measurement (*a*_*relMP*_ = *a*/*a*^*MP*^). The final odour response was than computed as the median response of all measured animals, in which sufficiently strong responses were observed (*a*^*MP*^ > 0.2‰).

To evaluate similarity of glomerular response spectra we computed the pairwise correlation distance, *d*_*r*_ (eq. 1):

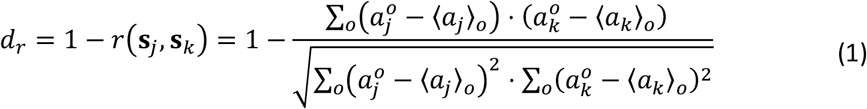

with 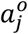 the response of glomerulus *j* to odorant *o*, and 〈*a*_*j*_〉_0_ the average response of glomerulus *j* to all odors. Thus, highly correlated response spectra will have distance zero, whereas fully anticorrelated spectra will have distance 2.

Based on this distance we obtained a hierarchical clustering using the unweighted pair group method with arithmetic mean (UPGMA) approach and evaluated clusters at different thresholds. We manually set thresholds to obtain clusters with distinct gaps between them. Thereby we restricted clusters to contain at least glomeruli of three different animals. To account for varying overall response strength across animals we normalized all responses to unit length response spectra 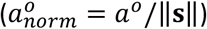.

### Physico-chemical characterization

For all measured molecules we obtained 3D structures via Pubchem (https://pubchem.ncbi.nlm.nih.gov/). We managed the structures using ChemAxon Instant JChem (version 5.9.4, 2012, http://www.chemaxon.com) and standardized them using the Clean3D function. For all molecules we computed 1600 descriptors of 19 descriptor blocks in eDragon (http://vcclab.org, ^37^), using all descriptor blocks except Information Indices. Furthermore we computed the EVA descriptors ^22,23^. To this end we optimized structures and computed vibrational frequencies in the GAUSSIAN software package using the B3LYP method and the 6-311G(d,p) basis set. Afterwards the vibration spectra were mapped to a continuous range [0,4000 cm^−1^] by placing a Gaussian kernel of unit height and varying width σ ∈ {1,5,10,20,50,100} on each frequency. The final descriptors sets 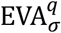 were then generated by sampling from this spectrum from *q* = 0 to *q* = 4000 in intervals of σ. We fitted activation models 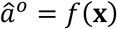 descriptor sets **x** to match the observed odour activations *a*^*o*^. We evaluated goodness of fit by calculating the coefficient of determination *R*^2^, eq. 2,

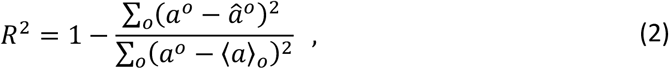

over all odorants *o*. Whereas this measure assessed how well the data could be described by a model, it does not reflect that sufficiently complex models could perfectly fit any functional relationship of the data without the ability to generalize to unseen data. Therefore we also evaluated the coefficient of determination for unseen data in a bootstrap approach ^38^. To this end, we fitted each model on 50 bootstrap samples *b*_*i*_ of the data and obtained predictions of odours 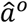 not contained in the bootstrap sample. We then calculated the predictive power as in eq. 3,

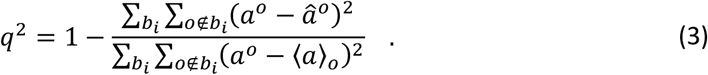

We evaluated the performance of univariate linear regressions, isotonic regression and unimodal isotonic regression ^39^ and multivariate Support Vector Regression (SVR) with Gaussian Kernel ^40^. All models were obtained via the scikit-learn module using default parameters ^41^. Scikit-learn was also employed to obtain the Multidimensional Scaling (MDS) of odour space distances. Prior of fitting SVR and MDS models we normalized all descriptors to zero mean and unit variance with respect to our in-house library of about 1000 odours.

We also employed normalized descriptors to obtain glomerular position in chemical space. To this end we assigned each glomerulus its barycentre of activation, eq. 4,

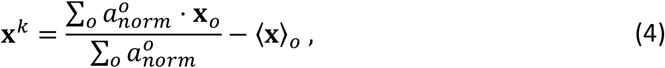

relative to the mean descriptor value 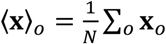 for all *N* odours in the measurement set. We then calculated the pairwise cosine distance of the barycentres (eq. 5),

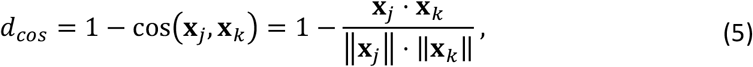

and obtained a hierarchical clustering thereof using the UPGMA approach.

### Replicability

We wish to encourage further use of the dataset and methods developed here. Therefore, we provide all code and data that is necessary to replicate each figure in this contribution. Please refer to the “Replication guide” in the supplementary material for links and instructions.

## Results

Our aim was to assess chemotopy in the dorsal olfactory bulb. We started by determining a detailed receptive range of the MOR18-2 glomerulus, from which we inferred a physicochemical model of the structure-activity-relationship for MOR18-2. The odorant profile and the physicochemical model enabled us to assess similarity of response profiles of glomeruli in the neighbourhood and evaluate the chemotopic order among glomeruli.

### The molecular receptive range of MOR18-2

We measured the response of MOR18-2 glomerulus by Intrinsic Optical Signal (IOS) imaging of the dorsal olfactory bulb (dOB, Fig. 1a) under stimulation with monomolecular odorants in MOR18-2-IGITL mice ^34^. The GFP label enabled direct localisation of the glomeruli on which MOR18-2 OSNs converged (Fig. 1b). Band-passing the signal reduced signal noise and separated the glomerular signal from the global unspecific signal ^42^ (Fig. 1c). Signals from individual glomeruli were then extracted using regularised Non-negative Matrix Factorisation (rNMF) (Fig. 1d-g, see methods and ^36^). The obtained glomerulus-like image components closely matched the original glomerulus outlines obtained from the GFP signal (Fig. 1e and f). This approach maximises the signal-to-noise ratio of the functional response as it maximises the area over which the glomerular signal is obtained. Fig. 1h depicts the spectrum of odorant responses in the component corresponding to MOR18-2 and the neighbouring component. As a result of the image processing method, there is little discernible signal spill-over between the two.

**Figure 1:**
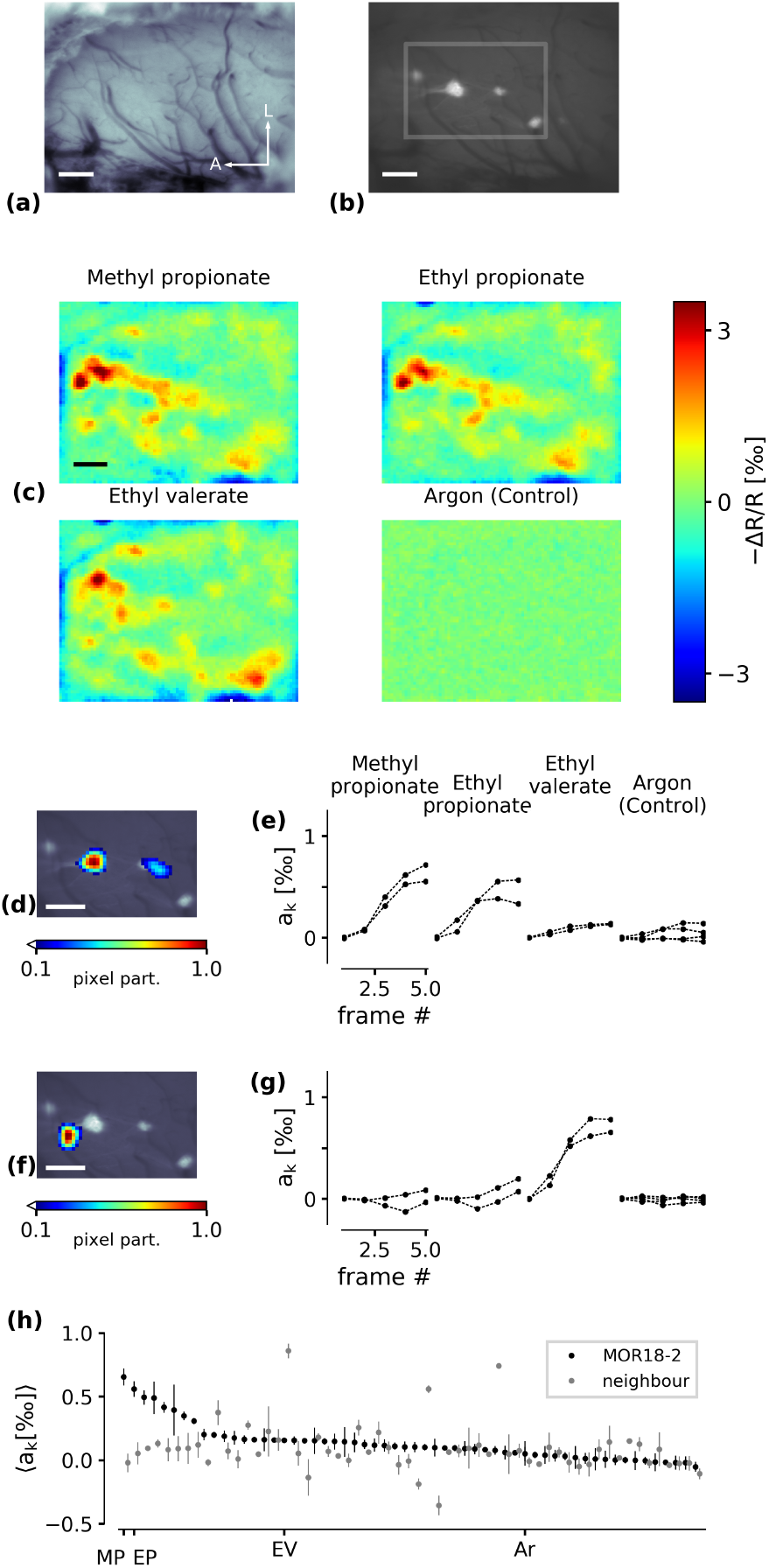
Automatic segmentation of putative glomeruli. (a) Image of the dorsal olfactory bulb preparation. A/L arrows denote the anterior/lateral orientation. Scale bar: 200 µm. (b) GFP image of MOR18-2 glomeruli. (c) Four examples of IOS odorant responses after preprocessing. (d) Pixel participation *x*_*k*_ of the rMF component that coincides with the location of the GFP signal. (e) Per-frame activation strength *a*_*k*_ of the component in (d) in response to the four stimuli depicted in (c). The odorant was released after frame 1. Argon (control) was repeated four times in this specimen, all other odorants were repeated twice. (f) A neighbouring component (putatively corresponding to a neighbouring glomerulus) and (g) the response to the four odorants. (h) Response spectra of the MOR18-2 mode (black) and the neighbouring mode (grey), average from two repetitions. Vertical lines denote minimal and maximal response. MP methyl propionate, EP - ethyl propionate, EV - ethyl valerate, Ar - Argon.

The IOS in the dOB reflects almost exclusively the activity of afferent olfactory sensory neurons ^43^, thus the functional response mainly conveys the response of OSNs expressing the MOR18-2 receptor.

In order to obtain an as complete as possible MRR of MOR18-2 glomeruli, we pursued an iterative screening approach, alternating between biological measurements and virtual screening ^44,45^. In each iteration, we built computational models based on the physicochemical properties of the biologically confirmed ligands. We used this model to screen odorant databases for potential ligands which we subsequently tested *in vivo*.

After discovering the first ligand (Ethyl acetate, compound 2 in Table ST1) by screening odours known to activate glomeruli in the dorsal bulb, we iteratively built hypotheses for further ligands based on the present set of revealed ligands. We acquired those hypotheses both empirically, by an intuitive notion of chemical similarity (e.g., chain length or bond saturation), and automated by virtue of virtual screening, using the computational models we built on the basis of physicochemical descriptors.

After several iterations of virtual/biological screening, we had obtained the response spectrum of the MOR18-2 glomerulus for 213 odorants from measurements in a total of 41 animals (Figure 2a). The response spectrum of MOR18-2 showed a rather narrow range of strongly activating odours. The structural diversity of the MOL18-2’s MRR is shown in Figure 2b, that depicts the most active ligands as well as some chemically similar but non-activating molecules. MOR18-2 was most sensitive to small aliphatic esters like Methyl Propionate, (compound 5), Ethyl Acetate (2), Propyl Acetate (4), Methyl Acetate (5), and the corresponding propionates (3,6,10). However, the most potent ligand we tested was Allyl Acetate (1), that has a double-bond in the terminal group. Interestingly, Isopropyl Acetate (9) yields relatively strong activity. It has a branched aliphatic group in the position of the double bond in Allyl Acetate. The structural variations of Allyl Acetate (1) and Isopropyl acetate (9) that we tested resulted in weaker responses, e.g. Allyl propionate (8) and γ-Isobutyl Acetate (30). Likewise, diverging from the central ester group led to lower-activity ligands, like Methoxyacetone (13), or Pyruvaldehyde (19). An unexpected observation was the strong response to Ethyl Aldehyde (7), as it is missing most of the features found in the other potent ligands. The full set of tested ligands, their measured activities and the *p*-value for difference to Argon can be found in supplementary table ST1.

**Figure 2:**
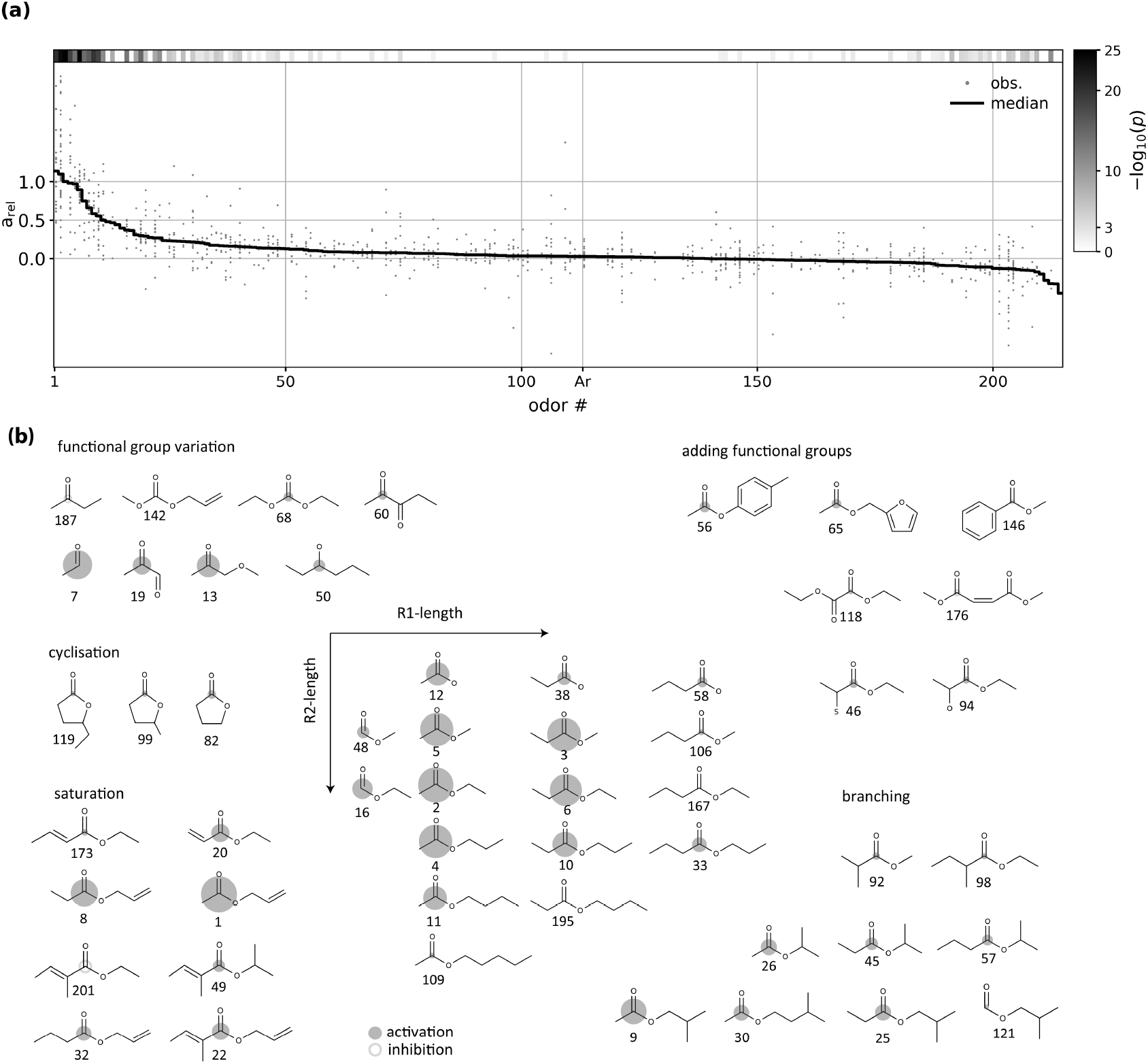
MOR18-2 response spectrum to 213 odorants. **(a)** All individual observed MOR18-2 response measurements (“obs.”, grey dots) and their median (black line). The column on the right indicates the *p*-value of a t-test that the measured response is different to control (Argon; Ar) measurements. See Supplementary Table ST1 for a list of all compounds, their response at MOR18-2, individual *p*-value and the number of replicates per odorant. **(b)** Molecular structure variation for some of the assessed odorants. Area of the circles is proportional to response strength. Molecules are numbered according to their MOR18-2 response rank (see Supplementary Table ST1 for ranks, names and CAS numbers).

We also observed some small, but statistically significant negative responses (e.g. compounds (187) and (201)). This observation may indicate that the glomerulus innervated by MOR18-2 receptor neurons shows inhibitory responses to these compounds. Since we assume that the intrinsic signal is dominated by ORN input, this observation implies that MOR18-2 normally emits a certain base firing rate that is decreased by these compounds. However, we cannot rule out the possibility that bandpass filtering in pre-processing induced artefacts that appear as inhibitory responses, for example through a very strong responses of a closely neighbouring glomerulus.

Glomeruli are not always arranged in a single plane in the rodent olfactory bulb. In order to exclude the effect of glomerular overlay an imaging method with 3D-resolution was therefore necessary. As an additional validation that the extracted IOS signals reflect mainly the glomerular input activation to the MOR18-2 glomerulus, we performed high resolution 2-photon synapto-pHluorin (spH) imaging in mice expressing both spH in all OSNs as well as GFP in MOR18-2 OSNs. To this end we first performed a 3D-anatomical scan of the resting spH fluorescence in which we manually outlined glomerular contours (Fig. 3a). We then performed functional spH imaging (Fig. 3b) and extracted glomerular activation via rNMF (Fig. 3d). Since the GFP-fluorescence was masked by the spH-fluorescence, we stained the olfactory bulb post mortem by Fast Red Violet (FRV). This way we made MOR18-2 glomeruli visible (Fig. 3c) and could assign them via anatomical landmarks (red arrows Fig. 3a/c) to the previously outlined glomerular contours.

**Figure 3:**
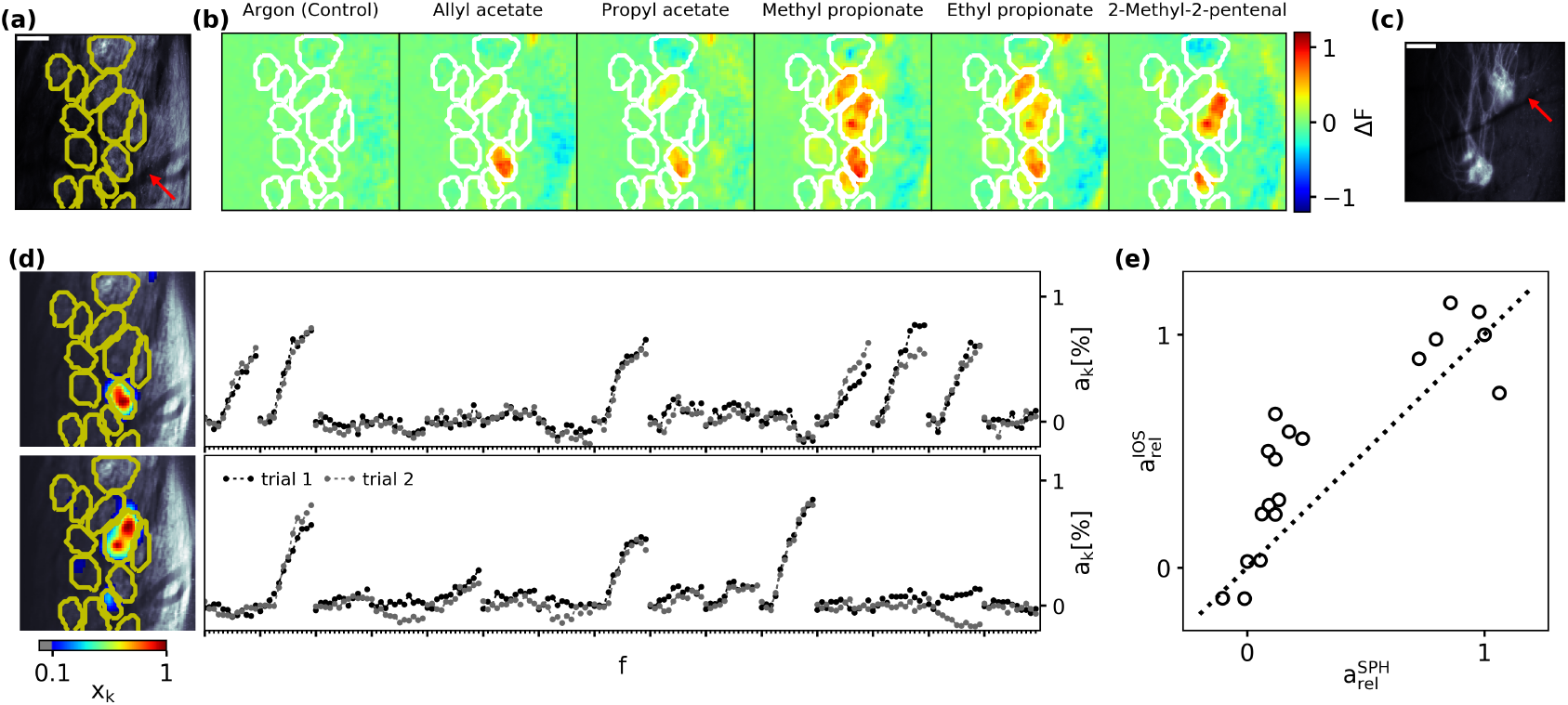
MOR18-2 SpH ligand spectrum. (a) Resting SpH-fluorescence in a patch of the dOB. We manually outlined glomeruli in the full 3D-stack (yellow contours). Anatomical landmarks (red arrow) were used to compare location to post mortem FRV images (Panel (c)). Scalebar 100µm. (b) Exemplary SpH odorant response maps. (c) Post mortem FRV image of MOR18-2 glomeruli. MOR18-2 location could be ascribed to the functional images via anatomical landmarks (red arrow). Scalebar 100µm. (d) Extracted activation of the MOR18-2 glomerulus component (top) and a neighbouring component (bottom). (e) IOS vs. SpH activation strength for all odorants.

The extracted spH activation of MOR18-2 glomeruli strongly affirmed the IOS derived ligand rank (Spearman’s rank correlation 0.92, *p* < 10^−7^). But in contrast to a rather continuously increasing response strength in the IOS spectrum, the spH-derived spectrum had a pronounced separation of strong and weak activations (Fig. 3e). This might be attributed to either a steep non-linear gain of spH fluorescence, or to a saturated IOS responses for strong activations, or a combination of both. In any case, the very high rank correlation between both spectra justified to rely on IOS measurements for subsequent analysis, which allowed us to record larger spatial extent and increased number of measured stimuli/repetitions than spH-imaging.

### Physico-chemical receptive range

Having obtained the MRR of MOR18-2, we asked if it could be described by a quantitative ligand-based model, that is, if response strength could be explained by a physico-chemical model of the odorants. To this end we first obtained 1600 physico-chemical descriptors via eDragon ^37^. These descriptors are subdivided into 19 blocks, ranging from simple scalar representations of molecules, like molecular weight or functional group counts, up to representations of the three-dimensional arrangement of properties (e.g. mass, charge) within the molecule. Additionally, we calculated the EVA descriptors in a range of spectral resolutions ^46^. We used EVA descriptors previously to determine structure-activity relationships for olfactory receptors ^23^. EVA descriptors have a free parameter, i.e. the width of the Gaussian kernel used for smoothing the vibrational line spectrum. We denote the particular variant used as EVA_x_, with x the kernel bandwidth in cm^−1^ (see methods for details).

We employed Support Vector Regression (SVR) models to explain and predict glomerular activation. Since we found that single descriptors are generally insufficient to describe structureactivity relationships in our data (see supplementary Results and Figure S3), we employed a multivariate approach. In a broad sense, SVR models estimate the response of the glomerulus to an odorant by forming a weighted sum of the responses to similar odorants, where similarity is determined in the underlying descriptor space. Therefore, for SVR models to become predictive, odorants evoking similar glomerular response must be proximal along some axis of the descriptor space. A multidimensional scaling (MDS) projection of odorants in the combined space of all eDragon and EVA_5_ descriptors showed that activating odorants clearly cluster together (Figure 4a). Consequently, the SVR model obtained in the combined eDragon/EVA_5_ space was able to capture the glomerular response to odorants in the training set (Fig. 4b), as well as generalise to odorants in the test set, which were not involved in model construction in any way (Fig. 4c).

**Figure 4:**
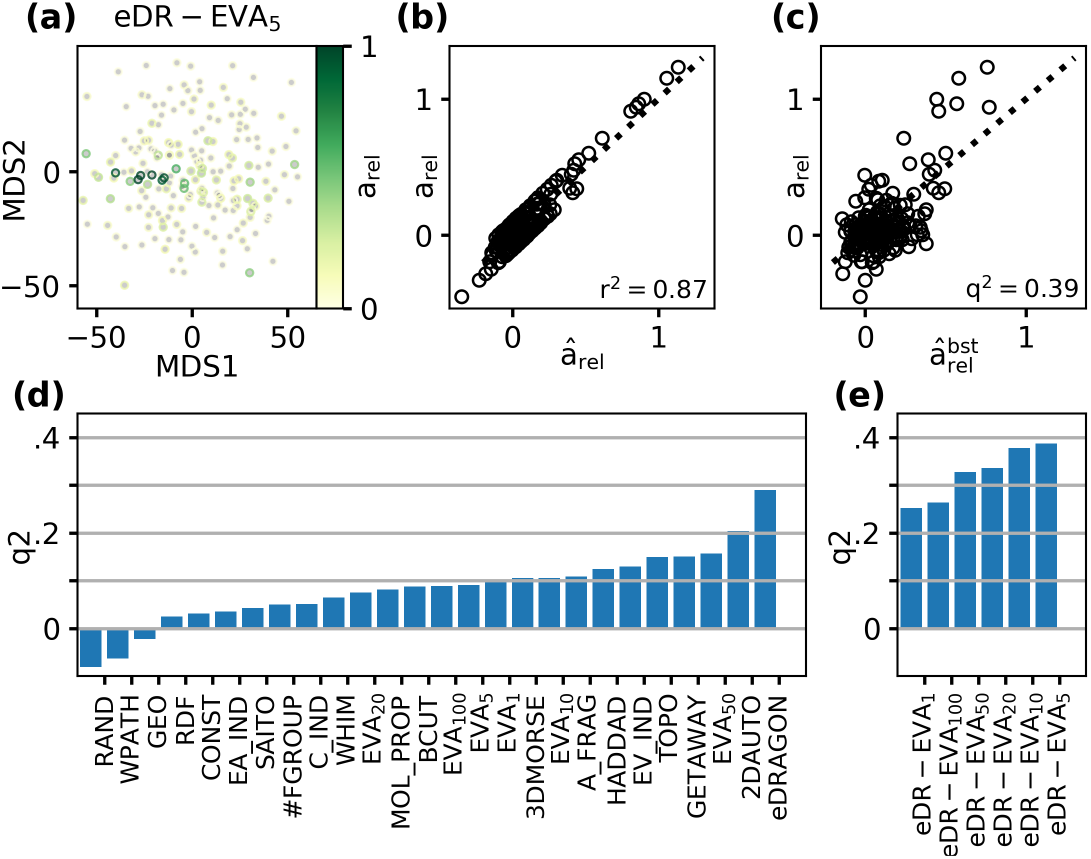
MOR18-2 MRR models. (a) 2D representation of odorant placement in eDR-EVA_5_ descriptor space via Multi-Dimensional Scaling (MDS). Circle edge colour depicts MOR18-2 activation of odour. (b) Observed MOR18-2 activation *a*_*rel*_ vs. model values 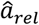 of a SVR regression models obtained in this descriptor space. The fraction of explained variance *r*^2^ is given in the lower right corner. (c) Observed activations *a*_*rel*_ vs. bootstrap model predictions 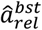 of the same model. The predictive power *q*^2^ is given in the lower right corner. (d) Predictive power *q*^2^ of SVR models in different descriptor spaces. (e) Same as (d) with combined eDragon and EVA descriptor spaces of different resolutions.

We evaluated which physicochemical descriptor sets would yield the highest predictive power in SVR models (Fig. 4d). We compared the full set of eDragon descriptors (eDRAGON), the logical subsets within them (see supplementary table ST2 for abbreviations), two subsets previously proposed to describe odour distances (HADDAD ^21^, SAITO ^20^ and the EVA descriptor at resolutions of 1, 5, 10, and 50 cm^−1^ (EVA_*x*_)). We found that the full set of eDragon descriptors yielded the best performance, followed by the 2DAUTO (2D-autocorrelations) and the odour-optimized HADDAD subsets. Notably, while the EVA descriptors did not outperform the HADDAD and SAITO descriptor subsets, they still yielded better predictions than most of the other eDragon subsets. We then tested whether predictive performance could further be enhanced by combining the eDragon descriptor set with the EVA descriptors (eDR-EVA_x_, Fig. 4e). And indeed, the combination of both sets improved performance by up to about 30 %, yielding models of MOR18-2 activation with a predictive power *q*^2^ ≈ 0.4. Thus we conclude that, despite its high dimensionality (2400 descriptors), proximity in the combined eDragon-EVA descriptor space provided the best estimate of chemical similarity of odorants with respect to their activation of MOR18-2.

### Tunotopic embedding

Since our imaging approach covered a considerable field of view across the dOB, it also included simultaneous recordings of glomeruli in the neighbourhood of MOR18-2, allowing to compare its MRR to other MRRs in the local glomerular ensemble. In general, the main class of MOR18-2 ligands (i.e., small esters) activated several glomeruli across the dOB (Fig. 1b and Supplementary Figure S1). Activity elicited by MOR18-2 ligands was therefore not restricted to a clearly defined sub-region. Likewise, the spatial location of the MOR18-2 glomeruli was not designated by any odorants which were solely activating this area. This raised the question about how the molecular response profile of the MOR18-2 glomerulus related to the tuning of its neighbours – that is, its *tunotopic embedding* in the dOB.

To investigate this tunotopic embedding we compiled a set of 48 odorants that we used to systematically probe the MRR of the glomeruli surrounding the MOR18-2 glomerulus (Table ST1, last column). This set contained most of the ligands that elicited a significant positive response in MOR18-2, along with some of their structural variations, therefore representing the previously determined MRR of MOR18-2. We measured the dOB IOS response in four GFP and three SpH mice and extracted glomerular response spectra (see methods). In the SpH mice, we validated the glomerular segmentation of the IOS via anatomical glomeruli outlines obtained in a 3D-scan of the resting SpH fluorescence (see ^36^). The GFP marker allowed us to identify the MOR18-2 glomeruli.

To investigate tunotopic relations between the extracted glomeruli, we calculated pairwise response similarity between all extracted glomeruli in all seven animals. Response similarity was assessed as the correlation distance *d*_*r*_ of the odour response spectra. Based on this distance we then obtained a hierarchical clustering of glomeruli (Fig. 5a). The spectra of the GFP-labelled MOR18-2 glomeruli displayed high mutual similarity, and they were all contained in a cluster with a low similarity threshold (red cluster in Fig. 5), suggesting that this approach is suitable to identify glomeruli corresponding to the same olfactory receptor across individuals.

**Figure 5:**
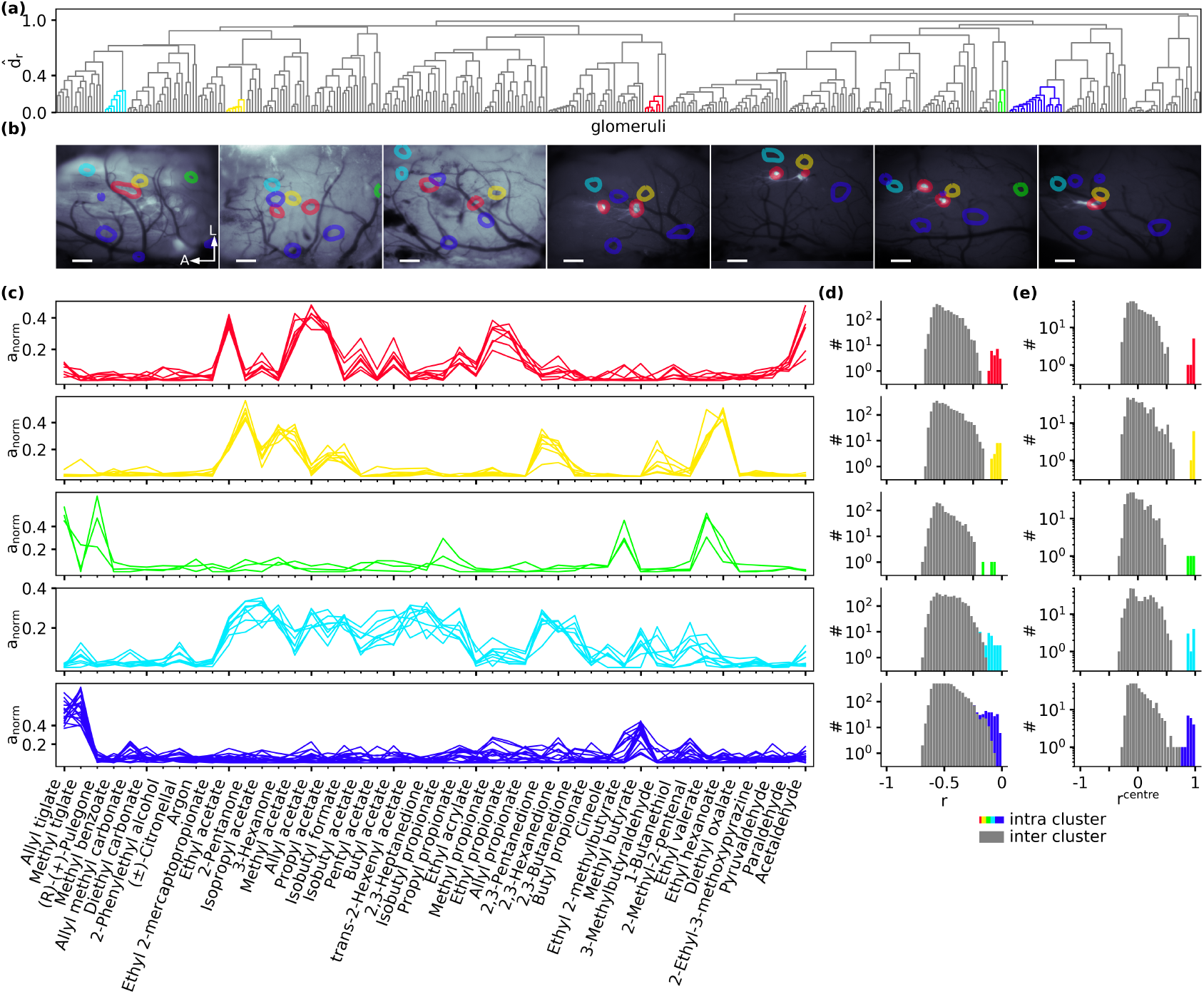
Functional identification. (a) Hierarchical clustering of correlation distance *d*_*r*_ between glomeruli of 7 mice. Length of branches depicts the average correlation distance 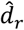 between clades. Properties of coloured clusters are shown in the following panels. (b) Spatial location of cluster members in the OBs of the seven mice in which the consensus odorant set was recorded. Colours according to cluster colouring in (a). Locations extracted from IOS imaging are overlaid on “green” images (first 3 animals) respectively GFP images (last 4 animals) indicating position of GFP labelled MOR18-2 glomerulus. Scalebars 200µm, A/L: Anterior/Lateral. (c) Odorant spectra of all glomeruli in each cluster. (d) Histogram of pairwise correlation *r* of all cluster members (coloured) and their correlation to remaining glomeruli (grey). (e) Histogram of correlation *r*^*centre*^ of all cluster members (coloured) and non cluster members (gray) to cluster prototype.

We observed several other low-threshold clusters that indicate a putative match of glomeruli across animals, indicated by the yellow, green, turquoise and blue clusters in Fig. 5. The glomeruli corresponding to those clusters were located at stereotypic positions across animals (Fig. 5b and Supplementary Figure S2). Their individual response spectra were highly overlapping (Fig. 5c). Histograms of pairwise similarity values of these glomeruli to all other glomeruli clearly showed that the putatively matching glomeruli shared high similarity to each other, while the similarity to non-members of the clusters was consistently lower, at least for the red, yellow and green clusters (Fig. 5d). For the turquoise and the blue clusters, the observed response spectra were not as clearly separated from the other glomeruli as for the red, green and yellow clusters. Nevertheless, a clear separation was possible when assessing the similarity to the cluster centroid, instead of a full pairwise comparison (Fig. 5e). In the case of the green cluster in Fig. 5, the cluster spanned only a subset of the animals – that is, the corresponding glomeruli could not be extracted in all 7 animals – probably, because it was located outside the recorded area in some of the animals. Nonetheless, we conclude that glomeruli with high mutual similarity likely correspond to glomeruli driven by the same class of olfactory receptor neurons, and that they can be identified by the similarity of their functional response spectra through hierarchical similarity clustering.

We next investigated the similarity response profiles of those putatively identified glomeruli to the MOR18-2 glomerulus. To this end we determined the correlation distance *r*_*MOR18-2*_ = *r*(*S*_*MOR18-2*_, *S*_*i*_) between the spectrum of a putative glomerular cluster *S*_*i*_ to the spectrum *S*_*MOR18-2*_ of the cluster corresponding to MOR18-2.

In Fig. 6a, clusters are highlighted if they exhibited a response correlation *r*_*MOR18-2*_ > 0.2. The hierarchical clustering procedure grouped four of those clusters together with the MOR18-2 cluster (coloured blue to violet in Fig. 6a), forming a ‘meta-cluster’. Closer inspection of their response spectra revealed that these glomeruli share an activation by propionates (Fig. 6c). These glomeruli did not only share similar activation but were also spatially proximal to each other on the dOB surface, forming a “patchy domain” around the MOR18-2 glomeruli (Fig. 6b).

**Figure 6:**
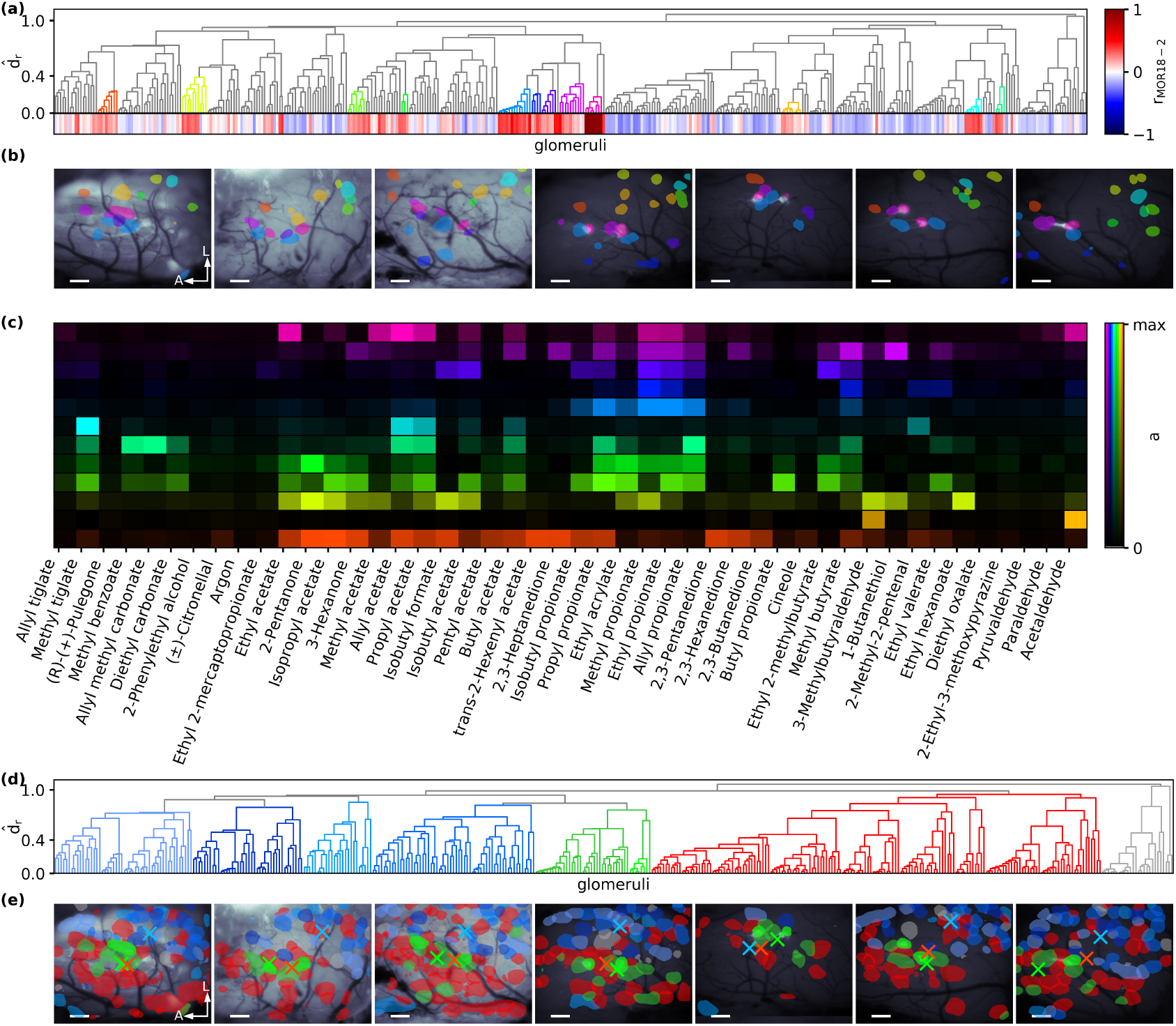
Tunotopic clustering of response spectra. (a) Hierarchical clustering of glomeruli of seven mice according to the correlation distance *d*_*r*_ between their response spectra. Each leaf corresponds to a glomerulus, and its correlation with the MOR18-2 cluster (pink) is colorcoded at the bottom of the panel. Clusters with *r*_*MOR18-2*_ >0.2 are coloured. (b) Spatial location of coloured cluster members in the OBs of all seven mice. Scalebars 200µm, A/L: Anterior/Lateral. (c) Median odour spectra of all clusters. Colours according (a). (d) Same hierarchical clustering as in (a) with colouring of “high-level” clusters. MOR18-2 is light-green. (e) Spatial location of cluster members in the OBs of all seven mice. Cluster centres of mass are marked by crosses. Colours according (d). Scalebars 200µm, A/L: Anterior/Lateral.

We quantified this observation by measuring the distance of all glomeruli to the nearest MOR18-2 instance, and comparing distances of patch members with those of non-patch members. We found that patch members are indeed more likely to be spatially proximal than non-patch members (supplementary Fig. S4A). Mann Mann-Whitney U statistics yielded a normalised U of 0.286 (*p*<10^−4^), i.e. when randomly drawing pairs from intra- and extra-patch glomeruli, the intra-patch glomerulus will be closer to MOR18-2 in 71.4% of the cases. A control analysis with 10,000 repeated shuffles of spatial distances showed no preference (supplementary Fig. S4B) and yielded an average normalised U_shuffled_ 0.51 (std. dev 0.05). It can be ruled out that the observed U has been obtained by chance (*p*=0, supplementary Fig. S4C).

Besides this local domain of glomeruli with mutually similar response spectra, MOR18-2’s spectrum was also correlated to other putative glomeruli located lateral-posterior to MOR18-2 glomeruli (orange, yellow, green and turquoise hues in Fig. 6b). However, each of those clusters was most similar to glomeruli uncorrelated to MOR18-2 - otherwise the clustering algorithm would have paired them up with MOR18-2. That is, although glomeruli with somewhat similar response profiles to MOR18-2 were located at distant locations, only those in proximity to MOR18-2 formed a meta-cluster of high mutual similarity.

We then asked whether the relative arrangement of such meta-clusters of spatially grouped glomeruli also followed a stereotypic rule that was conserved across animals. To this end, we chose a high similarity threshold to form clusters of glomeruli, obtaining essentially two ‘superclusters’ of glomeruli (Fig. 6d). The members of one supercluster were located predominantly in the lateral-posterior part of the dOB (blue glomeruli in Fig. 6e), while the other supercluster’s members were located mostly in the medial-anterior part (coloured red in Fig. 6d and e). The members of the MOR18-2 group (coloured green in Fig. 6e) are located in a region where those two ‘superclusters’ overlap. In 6 out of 7 animals, the axis defined by the centres of mass of the two superclusters aligned within 20°in the lateral-posterior to medial-anterior direction (supplementary Fig. S5A). A control analysis with shuffled glomerular identities (10,000 repetitions) yielded a bi-modal distribution of angles whose main axis did not align with the observed direction, except in one of seven animals. The observed distances between the centres of mass were larger than 95% of the distances obtained with shuffled glomeruli in six of seven animals (supplementary Fig. S5B).

In summary, we found that glomeruli were arranged at stereotypical positions regarding their response similarity to MOR18-2. Moreover, their spatial arrangement in the dOB correlated with the chemical response profile of the corresponding olfactory receptors. This holds on a local level, as a group of glomeruli with high overlap in their response spectra were arranged in a confined, but patchy, spatial domain around MOR18-2. Furthermore, we found that, on a larger scale, glomeruli in the dOB can be grouped in two ‘superclusters’ according to their response spectra, separated in a lateral-posterior, and a medial-anterior domain of response similarity.

### Chemotopic embedding

The tunotopic embedding of MOR18-2 that we observed suggested that there would exist, along some dimension of chemical space, also a chemotopic embedding, that would group glomeruli with a similar physico-chemical response profile profile together. Testing for chemotopy is, however, notoriously difficult since there is yet no clear-cut low-dimensional description of odorant space that would allow to define a sensory topology of the like that is found in visual or somatosensory systems. Therefore, investigating a potential chemotopic embedding required defining a chemical space (i.e., chemical similarity) in which odorants are arranged, and to define the position of glomeruli within this space.

As chemical space we choose the combined eDR-EVA_5_ space since it proved best to characterize the MRR of MOR18-2 (see section MOR18-2 physico-chemical receptive range). Furthermore, we placed the origin of the coordinate system at the mean descriptor value ⟨**x**⟩_*o*_ of our odour set. The location of a glomerulus *k* within this space was defined by its barycentre (i.e., the centre of mass) **x**^*k*^ with regard to its activation spectrum (see eq. 4 in the Methods section). Hence, a glomerulus responding with equal activity to all odorants would be positioned at the origin of chemical space.

We then obtained the similarity of glomeruli to each other by calculating their cosine distance *d*_*cos*_ = **x**^*k*^ ⋅ **x**^*l*^, which is a measure of the angle between the glomerular positions (see eq. 5 in the Methods section). In other words, glomeruli were considered similar, if their barycentres of activation were located in a similar direction from the origin.

Based on these pairwise distances we again obtained a hierarchical clustering (Fig. 7a). All MOR18-2 glomeruli were grouped together in a low threshold cluster (violet cluster in Fig. 7a & b), similar as with tunotopic clustering (cf. red cluster in Fig. 6a & b). However, in contrast to the tunotopic case, the MOR18-2 glomeruli were not embedded in a small scale local chemotopic domain, but only in a meta-cluster of lateral-posterior located glomeruli (violet-blueish cluster in Fig. 7c & d). This meta-cluster was contrasted by an adjacent meta-cluster of medial-anterior glomeruli (orange and yellow cluster in Fig. 7c & d). But on the topmost level of clustering more distant glomeruli were widely dispersed in both domains (grey glomeruli in Fig. 7c & d). In summary, our approach could reveal a clear chemotopic arrangement of glomeruli with high similarity to MOR18-2. However, the observed chemotopic structure was less obvious than when assessing response similarity on a tunotopic basis.

**Figure 7:**
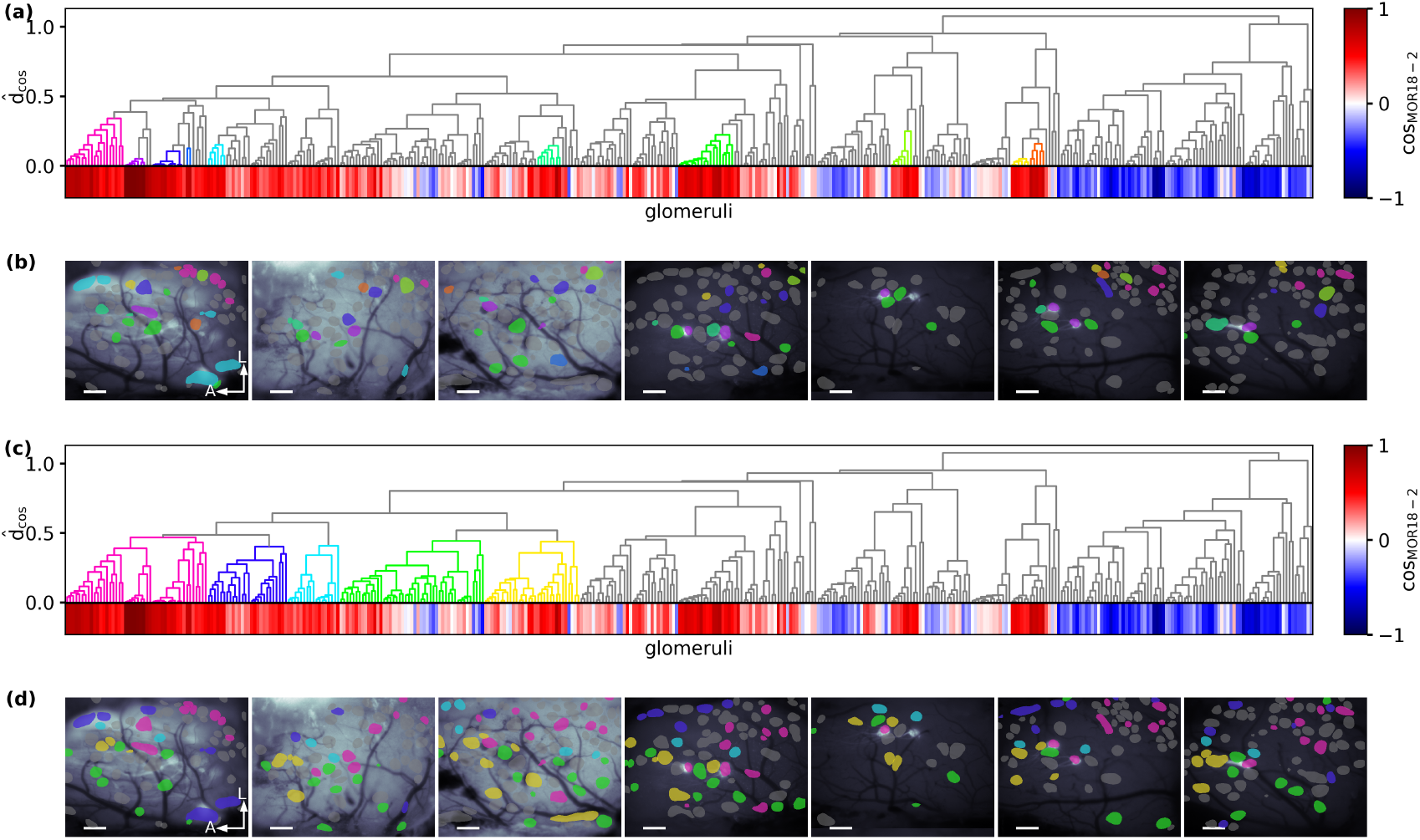
Chemotopic clustering of response spectra. (a) Hierarchical clustering of glomeruli according to the cosine distance *d*_*cos*_ between their chemical response spectra from the seven animals in which the consensus odorant set was recorded. For each glomerulus *k* its angle cos(*x*_*MOR18-2*_, *x*_*k*_) with the MOR18-2 cluster is color-coded below the dendrogram. Several representative clusters that share some chemical similarity with the MOR18-2 cluster (blue) are coloured. (b) Spatial location of cluster members in the OBs of all seven mice. Coloured locations correspond to representative clusters selected in (a). Scalebar 200µm, A/L: anterior/lateral. (c) Same hierarchical clustering as in (a) with colouring of meta level clusters at close distance to the MOR18-2 cluster. (d) Spatial location of cluster members in the OBs of all seven mice. Colours according (c). Scalebar 200µm, A/L: anterior/lateral.

Whilst the chemotopic clustering failed to reveal a local “patch” as observed for tunotopy, we nevertheless found that glomeruli most similar to MOR18-2 were predominantly located in its immediate vicinity. In order to quantify the relationship between spatial proximity and similarity of chemical receptive fields of glomeruli relative to MOR18-2, we divided glomeruli into “near” and “far” glomeruli according to a spatial distance threshold, and tested for the “near” glomeruli being chemically more similar to MOR18-2 using Mann-Whitney U statistics. The test is always significant (*p*<0.05) up to a distance value of 180µm, and the normalised U values (i.e., the odds of far glomeruli being more chemically similar than near ones in pairwise comparisons) were less than 0.5 for all distance thresholds p to 490 µm (supplementary Fig. S6A). Controls with shuffled chemical and spatial distances yielded normalised U values around 0.5 (*p>*0.05) for all tested distance thresholds (supplementary Fig. S6B and C). This analysis confirmed our qualitative observation that the receptive fields of glomeruli in spatial proximity to MOR18-2 are more likely to chemically similar to MOR18-2’s MRR.

## Discussion

In our study, we profiled in detail the MRR of MOR18-2 glomeruli and their embedding in the glomerular code. MOR18-2 glomeruli mainly responded to small esters and locally embedded in a patchy domain of glomeruli sensitive to short chain propionates. Furthermore, we could determine a physico-chemical receptive range of MOR18-2 glomeruli within the multidimensional eDR-EVA_5_ space via SVR models. We also observed a chemotopic representation of this space within the dorsal OB, albeit less obvious than the tunotopic arrangement.

### MOR18-2 receptive range

We aimed to obtain the MRR of MOR18-2 as completely as possible, but it is of course infeasible to screen all existent odours. Our informed search for candidate ligands was based on predictive physico-chemical models, but also (to a lesser amount) manually selected search candidates. In spite of this directed search, it is unlikely that our set of ligands exhaustively represented the full response repertoire of MOR18-2. Indeed, our predictive SVR model, as ascertained on bootstrap samples, suggests further possible ligands, which could be evaluated in future screening campaigns. To our knowledge this study makes MOR18-2 one of the best explored olfactory receptors ^47^. Based on the data published along with this report, it will be possible to construct new predictive models, potentially including any ligands discovered later.

Recently, acetic and propionic acid have been reported as ligands of MOR18-2 receptors expressed in the kidney ^48^. In contrast, our results rather identified the corresponding esters as ligands. Although their study did not contain any of the ligands revealed in our study, there could be systemic reasons for such a deviation. For example, we presented odour stimuli in the gaseous phase, taken from the headspace of a solution, while Pluznick et al. applied the liquid phase. Since acids tend to have much lower vapour pressure, the effective concentration that reached the receptors might have been considerably lower for acids than for the corresponding esters. Another potential reason for this discrepancy is the potential conversion of esters into the corresponding alcohols and acids by enzymes in the mucus ^49^.

Assuming that esters are generally converted to their corresponding acids shines an interesting light on our results. The best 12 ligands we identified are all esters of either acetic or propionic acid. Hence, all of their responses could be explained by the receptor responding to either of those acids. Asserting whether this is indeed the case would require administering esterase blockers in the mucosa prior to the experiment, which is beyond the scope of the current study.

As our aim was to obtain an as complete MRR of MOR18-2 as possible, we did not obtain doseresponse curves for odorants, since obtaining those curves would have dramatically reduced the amount of odorants to be measured. Instead, all odorants have been used at very high concentrations – liquid odorants in highest purity available and solid ones in saturated solutions. We are therefore unable to make any statement of concentration effects on the MRR.

### Physico-Chemical Receptive Range

In this study we did not only determine the MRR of MOR18-2 but also translated it via SVR models to a physico-chemical receptive range in the eDR-EVA_5_ space. We carefully validated our predictive models using bootstrapping. Previously, we successfully used a very similar approach for Drosophila ORs ^23^, so we are confident that physico-chemical range we obtained is a reasonably good estimate of the true range. Naturally, an ultimate confirmation of further model predictions would require successful biological screening — which in turn would enable improving the predictive models, and so on, continuing the iterative approach to virtual/biological screening that we employed here.

We obtained the best results in describing the physico-chemical receptive range by combining features based on molecular vibrations (EVA) with features containing descriptions of the molecular shape (eDragon). Note that this provides neither evidence for nor against the highly disputed theory of vibrational olfaction ^50^. In our model, molecular vibrations are considered as a kind of “fingerprint” of the molecular graph, rather than a mechanistic interpretation of molecule-receptor interaction.

### Tunotopic domains

Our results provided evidence for tunotopic domains at different scales: two global domains, a lateral-posterior and a medial-anterior one, and a local domain around the MOR18-2 glomeruli. These domains qualitatively resembled the domains described by Ma et al. (2012), characterised by a global partitioning and some local domains, although the domains we found had a slightly different local layout. The large-scale partitioning of the dOB into two tunotopic medial-anterior and lateral-posterior top-level domains has been described in a similar fashion across different studies. It resembles the domain separation of class 1 (medial-anterior) and class 2 receptors (lateral-posterior) ^51^. It is also in line with the MOR18-2 glomeruli being located close to the border of class 1 and class 2 domains ^52^. In contrast, the local tunotopic clusters we identified diverge from those identified in other studies, probably because of the different odorant sets that were used to identify tunotopic groups of glomeruli.

Interestingly, despite being a class 1 receptor, MOR18-2 is tunotopically embedded in the lateral-posterior domain which is presumably composed to a large extent of class 2 receptors. This could indicate that, albeit its physiological separation ^52^ the functional transition between receptor class domains is rather continuous.

Mori and colleagues ^14,51^ also observed a compartmentalisation of the dOB into local response clusters. They attributed long chain ester responses to cluster A (within class 1 receptor domain) and cluster B (within class 2 receptor domain) but did not attribute any cluster to small esters. However, their cluster assignments leave undesignated space especially at the border of domain 1 and 2 where we observed the propionate responding cluster of MOR18-2. This area is potentially populated with TAAR glomeruli ^53^.

Taken together, our findings confirm the hypothesis of a tunotopic layout of the olfactory bulb. This was disputed by ^17^ who observed “that nearby glomeruli were almost as diverse in their odorant sensitivity as distant ones”. In their study, they evaluated the dependence of glomerular spectrum similarity with regard to their spatial distance and did not find strong effects. A potential reason for the different result could be that a highly diverse odour set such as the one used on that study is not suited to resolve the similarity in MRRs among local response clusters. For example, the local response cluster of MOR18-2 would not emerge using an odour set containing only acetates, but lacking the corresponding propionates (which differ only by one additional Carbon atom).

### Chemotopic domains

Besides tunotopic domains, we did also observe a locally confined cluster of glomeruli with chemical similar response spectra. This chemotopic organization was less pronounced than the tunotopic organization, for several potential reasons. First, since there is no canonical definition of chemical similarity, any relation between chemical similarity and spatial proximity is not canonically defined, either. Our definition by the eDR-EVA_5_ descriptor space is suitable to describe the MRR of MOR18-2, but it might still be too imprecise to reveal physico-chemical structure-activity relationships in local chemotopic domains. Second, since our odour set was constructed to contain MOR18-2 ligands, it naturally would not cover the full ligand spectrum of other receptors. While we made this choice deliberately in order to resolve the local domain (in a chemical sense) of MOR18-2, our set of odorants will not be optimal to describe the MRRs of its neighbours. Since we represent the chemical receptive field of all glomeruli as a weighted combination of their response to the measured odorants, a full comparison of those receptive fields would require the key odorants of neighbouring glomeruli also being present in the set. This does not so much pose a problem when assessing tunotopic distance, since the response spectrum correlations are only influenced by the identity, but not the chemical properties of the ligands. In contrast, chemotopic distance is all about the chemical properties of the shared and unshared ligands. A complete assessment of chemical similarity between MRRs can, in theory, only be assessed if both MRRs are represented equally well with a number of ligands that sufficiently represent the region of chemical space in which those MRRs are located. Future research in this area should focus on identifying the MRR of neighbouring glomeruli to a much higher extent than in the present study, thus enabling better resolution of chemical similarity between MRRs.

A rigorous quantitative assessment of chemotopy is further complicated by the ambiguity of glomerular locations on the dOB surface. Most animals tested had multiple instances of the MOR18-2 glomerulus that were in the same area, but not immediately adjacent. In that case, it is impossible to provide a distance metric that appropriately captures chemical distances between MOR18-2 glomeruli and any other identified glomerulus. Possible approaches to deal with this ambiguity include using the centre of gravity of MOR18-2 locations as a reference, or running separate assessments for all instances of MOR18-2 glomeruli. Both approaches will result in non-zero distance between glomerulus instances that should be treated as identical for the purpose of mapping, violating the definition of a distance metric, compromising the rigour of a quantitative analysis based on such approximations. We partly circumvented this complication by adopting pairwise comparisons of glomerular locations in quantitative assessments, but the principal problem remains.

To conclude, we investigated the olfactory bulb’s topographic layout with regard to an extensively characterized MRR of a single glomerulus. This provided further evidence for a functional topographic layout according to glomerular response profile similarity. Nonetheless also this approach still suffered from the incomplete representation of MRRs. Therefore, it once more emphasises the demand of a full characterization of olfactory MRRs.

## Supporting information

Supplemental results, figures, and tables

## Additional information

### Funding and Acknowledgements

This works has been enabled by the Priority Programme SPP1392 “Integrative Analysis of Olfaction” funded by Deutsche Forschungsgemeinschaft (DFG), under grant nos. SCHM2474/1-1 & SCHM2474/1-2 (MS) and SP1134/1-1 & SP1134/2-1 (SPP 1392) (HS). Additional funding was provided by DFG under grant no. FOR 643 (HS). We are deeply grateful to Prof. Giovanni C Galizia and for creating a sense of community among olfaction researchers in Germany, and all members of SPP1392 for inspiring discussions.

### Author contributions

MS & HS designed the project and acquired funding. HS and JS1 (Jan Schumacher) designed and performed biological experiments. JS2 (Jan Soelter) designed and carried out the data analysis, with supporting analysis by MS, HS and JS1. MS and JS2 curated the data analysis code for easy reproducibility and made it available on github. JS2 and MS wrote the manuscript, with support from HS. All authors reviewed the manuscript.

### Data availability statement

All code and data to replicate all figures in this manuscript and the supplement are made available under an open source license. Code (in the form of jupyter notebooks) is available at https://github.com/Huitzilo/glomcentric_code/releases/tag/2.0 and https://doi.org/10.5281/zenodo.1745464; data is available at https://doi.org/10.5281/zenodo.1297376.

### Preprint

A preprint of this work has been deposited on *bioRxiv*, doi: 10.1101/489666 ^54^.

