## Supplemental results, figures, and tables for "Computational exploration of molecular receptive fields in the olfactory bulb reveals a glomerulus-centric chemical map"

#### Replication guide

We provide the complete codebase and data to replicate all figures in this contribution. Jupyter notebooks for replicating figures are provided at https://github.com/Huitzilo/glomcentric_code. The reference release is v2.0, archived at https://doi.org/10.5281/zenodo.1745464. Data can be found at https://doi.org/10.5281/zenodo.1297376 .

Steps to replicate:

1. Download and extract the data (5 GB download, 13 GB extracted).
2. Clone the code repository (will need about 130 MB after replication).
3. Required python packages and versions are listed in soelteretal.yml in the code repository. This file can be imported into Anaconda Python (python 2.7). Set up and activate a virtual environment (or conda environment) with the exact versions provided. We strongly encourage using the exact versions provided, since version conflicts are likely to cause errors that are difficult to detect and diagnose.
4. Launch jupyter notebook in the top-level directory of the code repository.
5. Launch MOB_Fig_decomp_and_spec.ipynb and follow the instructions.

Use the following jupyter notebooks to reproduce each figure:

Figures 1 and 2: MOB_Fig_decomp_and_spec.ipynb

Figure 3: MOB_Fig_SpH.ipynb

Figure 4: MOB_Fig_univar_multivar.ipynb

Figures 5 and 6: MOB_Fig_tunotopy.ipynb

Figure 7: MOB_Fig_chemotopic_clustering.ipynb

Reporting comments, bug reports and other issues is encouraged and should happen via the "issues" functionality in the github repository.

#### Supplemental results: Single descriptor models

We tested if a single descriptor is sufficient to explain the receptor activation. That is for each descriptor we evaluated different functional relationships between its value and receptor activation:

I. Linear increase of activation with descriptor value (linear regression)

II. Arbitrary (non-parametric) monotonic increase of activation (isotonic regression), e.g. saturated and/or thresholded increase of activation with descriptor value

III. Peak activation at intermediate descriptor value with monotonic decreasing activation both to lower and higher descriptor values (unimodal regression). This could for example be a scenario for chain length dependency.

For each of the three model classes Figure S3a depicts the best model over all descriptors with respect to the coefficient of determination $R^2$, i.e., the fraction of explained variance. With increasing model complexity more and more of the response variance became explained (Fig. S3b). However, none of the models could correctly capture the response to all odours. This shortcoming was especially apparent when evaluating the predictive power of the models (Fig. S3c). To this end we fitted the models to bootstrap sub-samples of the data. We then compared predictions to observed values for odours excluded in the sub-sample and calculated the coefficient of determination $q^2$ for unseen odours (see methods). The predictions both included odours with erroneous assigned activation and missed activation. This was reflected by a coefficient of determination $q^2$ below 0.25.

#### Supplemental figures

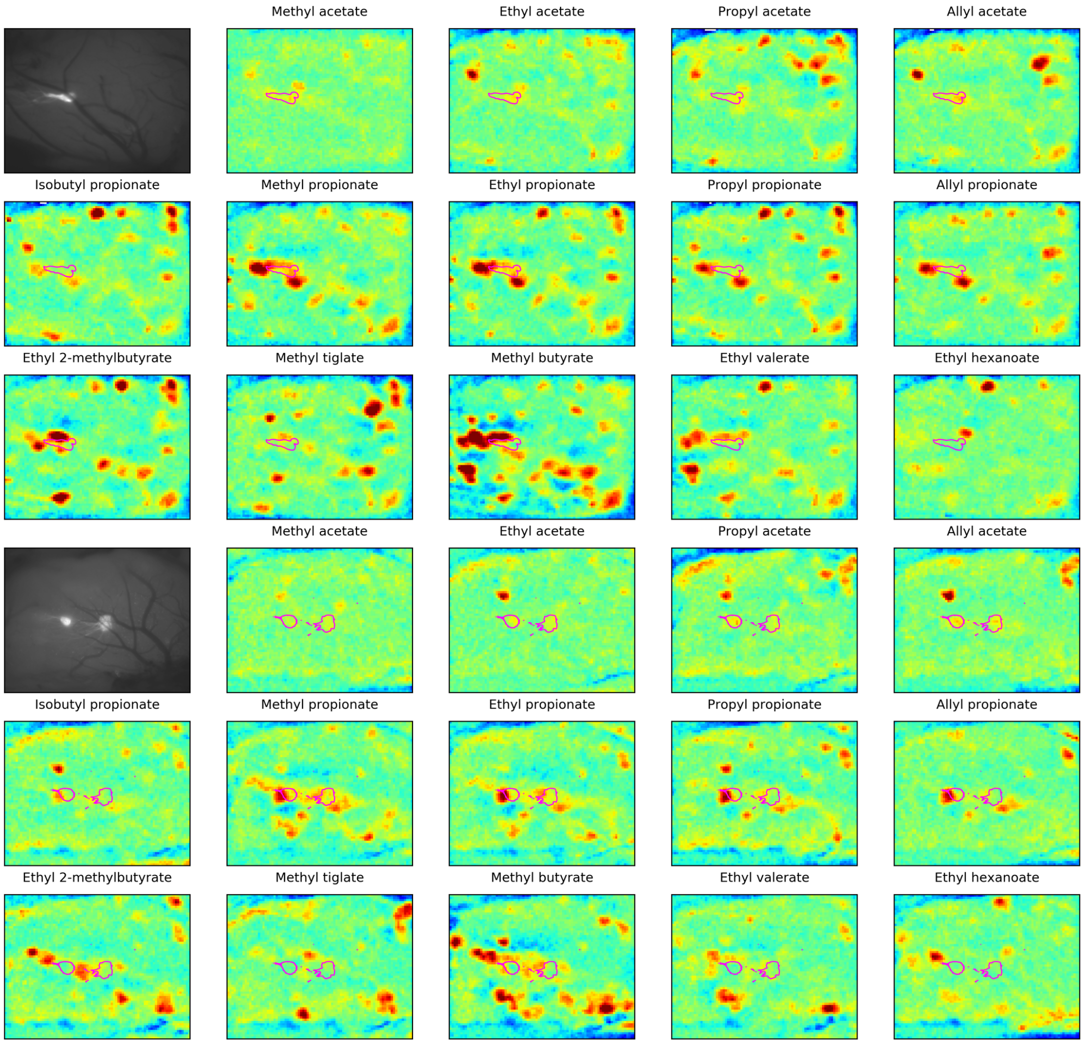


**Figure S1: Exemplary Odorant maps**. Odorant response in the dOB for a selection of small ester in two different animals. Position of MOR18-2 is marked by red contour lines.

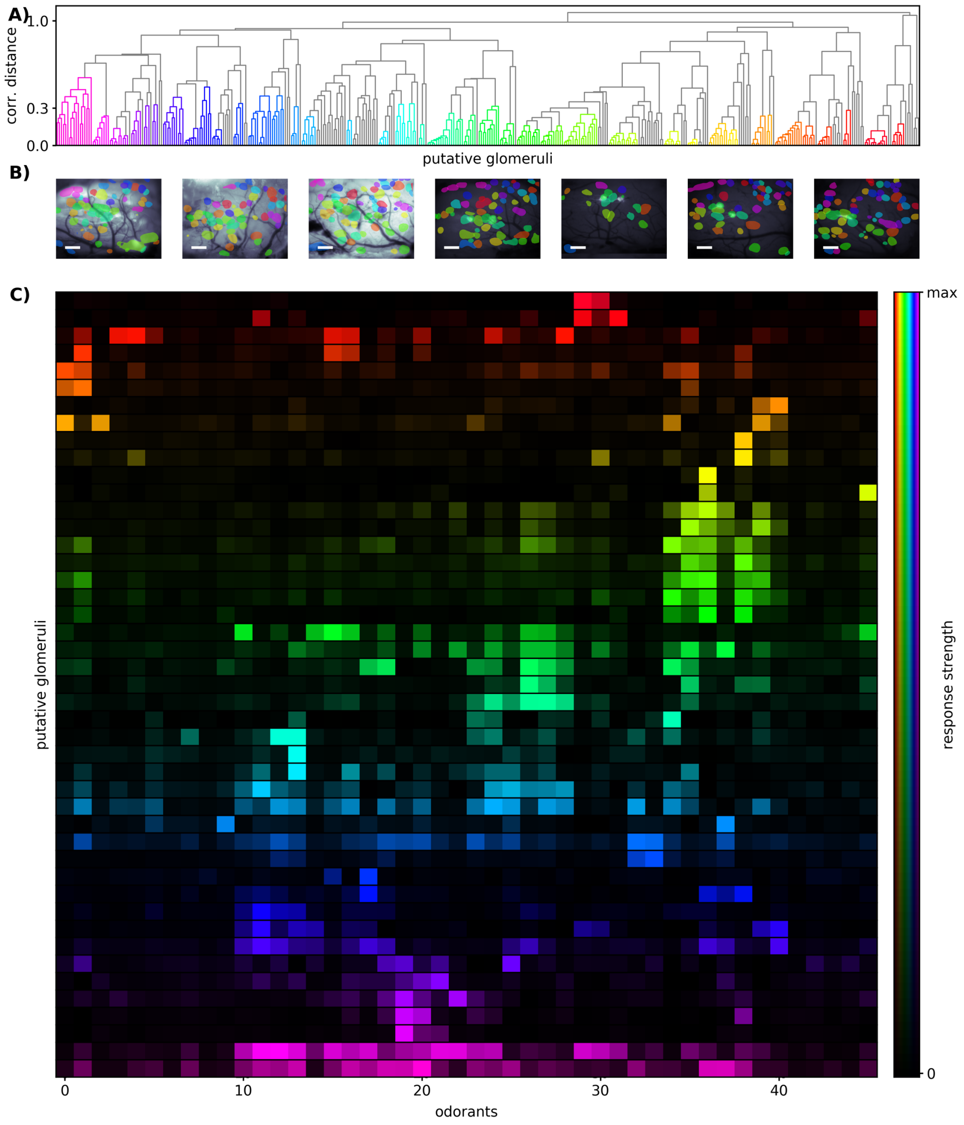


**Figure S2: Response Clustering** (a) Hierarchical clustering of correlation distance $d_r$ between glomeruli of 7 mice. Length of branches depicts the average correlation distance $\hat{d}_r$ between clades. Properties of coloured clusters are shown in the following panels. (b) Spatial location of cluster members in the OBs of all seven mice. Colours according to cluster colouring in (a). Locations extracted from IOS imaging are overlayed on green images (first 3 animals) respectively GFP images (last 4 animals). (c) Median odour spectra of all glomeruli in each cluster.

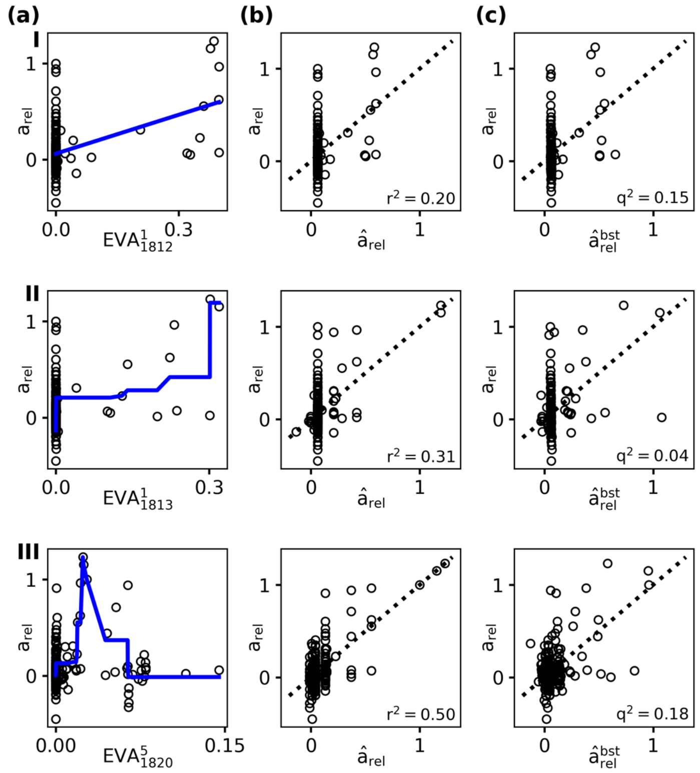


**Figure S3: Single-descriptor models.** (a) MOR18-2 activation $a_{rel}$ in dependence of descriptor values and corresponding regression models (blue lines): I. linear regression, II. isotonic regression and III. unimodal regression. Each descriptor was chosen to represent the best possible regression model. (b) Observed activations $a_{rel}$ vs. model values $\hat{a}_{rel}$ of the regression models depicted in (a). Resulting fractions of explained variance $r^2$ is given in the lower right corners. (c) Observed activations $a_{rel}$ vs. bootstrap model predictions $\hat{a}_{rel}^{bst}$ of the same models. Resulting predictive power $q^2$ is given in the lower right corners.

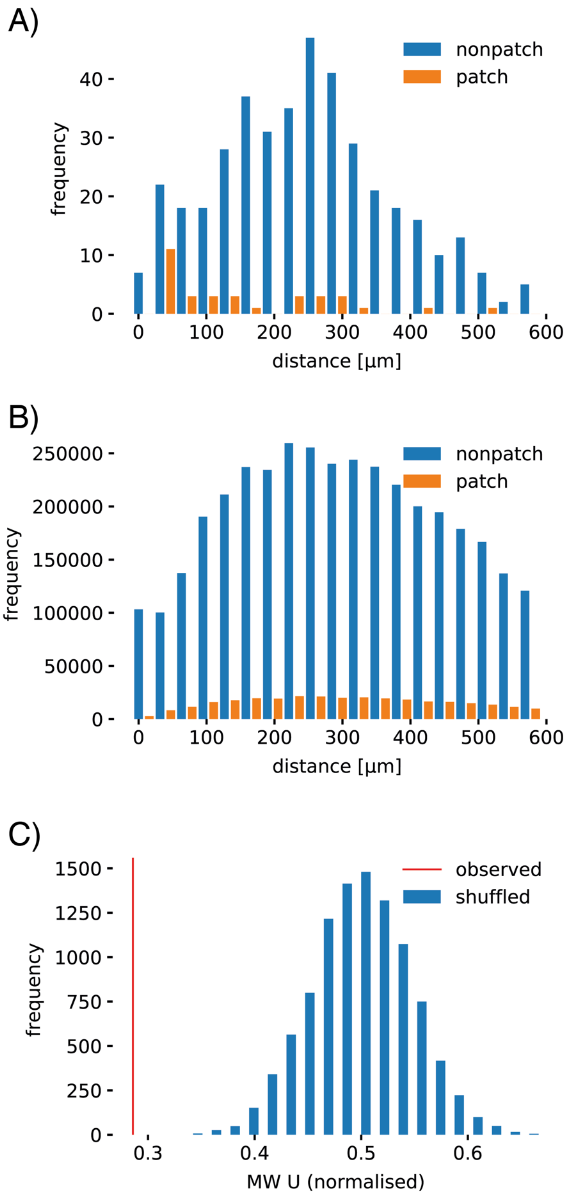


**Figure S4:** A) Histogram of pairwise distances between glomeruli that belong to the "patch" observed in hierarchical clustering, vs. those that do not belong to the ("non-patch"). Mann-Whitney U statistics yield a normalised U of 0.286 ($p<10^{-4}$). B) Controls with glomerular locations shuffled (10,000 repetitions). C) Histogram of Mann-Whitney U statistics; normalised, i.e. odds of non-patch patch glomeruli being more proximal than patch glomeruli in pairwise comparisons over 10,000 shuffled repetitions. The observed U value is smaller than 100% of the shuffled controls, therefore it can be ruled out that the observed U is obtained by chance ($p$=0).

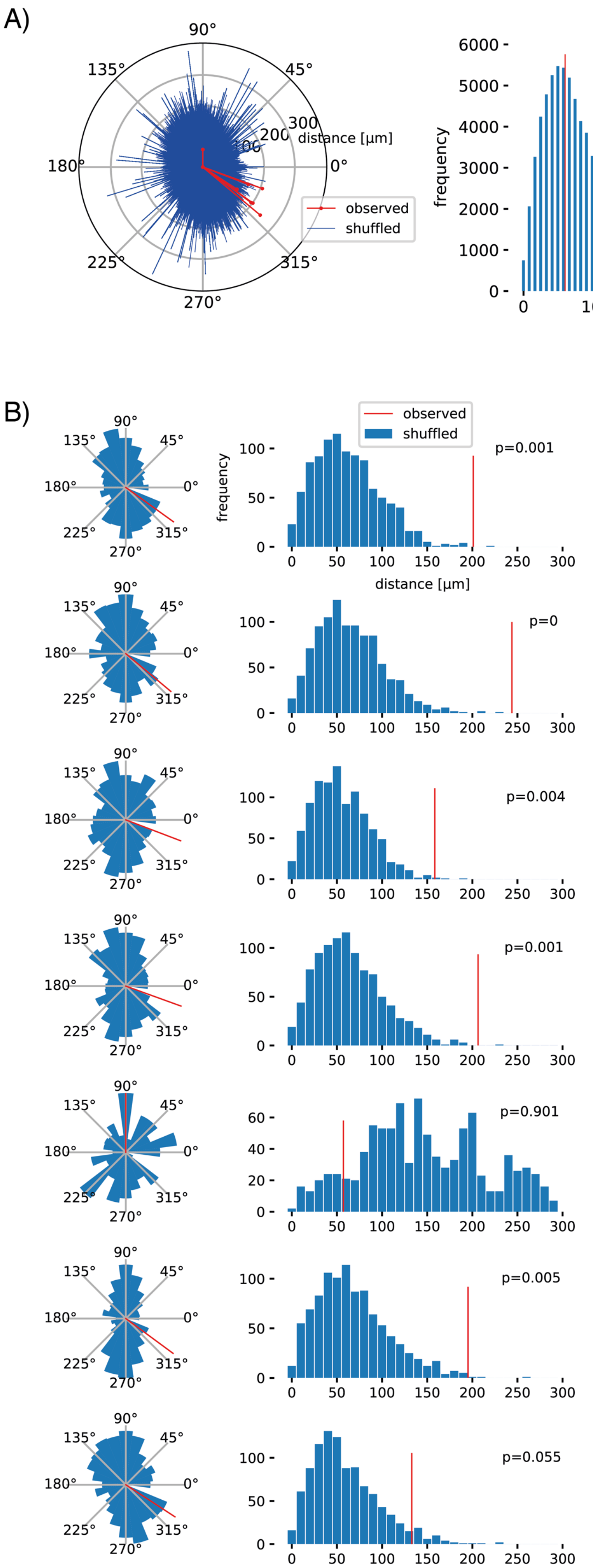


**Figure S5:** A) Left: Polar plot of observed vectors between centres of mass of red and blue meta-clusters (red), and 10000 shuffled controls (blue). Right: Histogram of distances between the cluster centres as obtained from 10,000 shuffled controls. Red lines indicate observed values. B) Per-animal polar histograms of directions of vectors between centres of mass (left) and distance histograms (right) for 10,000 shuffles. Observed angles and distances are indicated in red.

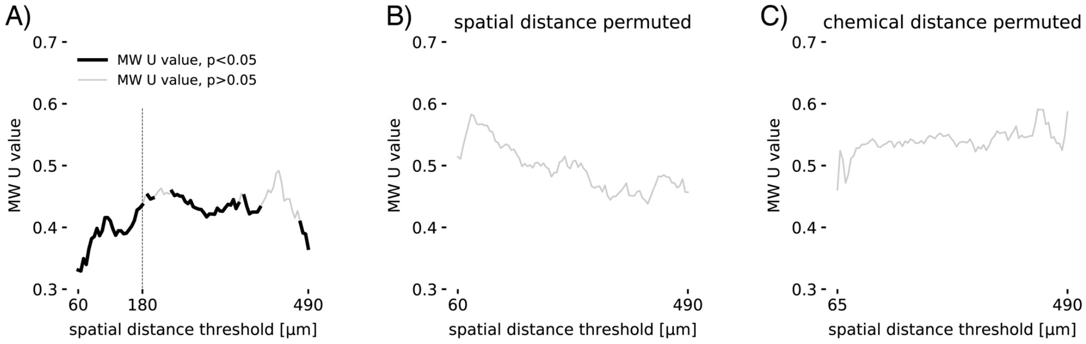


**Figure S6:** A) Normalised Mann-Whitney U statistics for chemotopic distance of receptive field centres being closer to MOR18-2 within a distance radius or outside. Normalised U values are always below 0.5 and significant ($p<0.05$) for radii up to 180 µm. B) U statistics for shuffled spatial distances, C) shuffled chemical distances. U values for shuffled controls are close to 0.5 and $p>0.05$ for all radii.

### Supplementary tables

**Supplementary Table ST1:** List of all measured odours ranked according to their response at MOR18-2. *P*-value indicates t-test outcome for difference from Argon response, $\log_{10}(p)$ is plotted in Fig. 2.

| Index | CAS | Name | MOR18-2 response | *p*-value | $\log_{10}(p)$ | in Fig. 3 | in Fingerprint/ Fig. 5/6 | Num. repl. |
|---|---|---|---|---|---|---|---|---|
| 1 | 591-87-7 | Allyl acetate | 1.14 | 0.0000 | 20.40 | yes | yes | 19 |
| 2 | 141-78-6 | Ethyl acetate | 1.10 | 0.0000 | 23.89 | yes | yes | 46 |
| 3 | 554-12-1 | Methyl propionate | 1.00 | 0.0000 | 90.28 | yes | yes | 46 |
| 4 | 109-60-4 | Propyl acetate | 0.98 | 0.0000 | 18.49 | yes | yes | 17 |
| 5 | 79-20-9 | Methyl acetate | 0.97 | 0.0000 | 13.83 | yes | yes | 8 |
| 6 | 105-37-3 | Ethyl propionate | 0.90 | 0.0000 | 34.11 | yes | yes | 18 |
| 7 | 75-07-0 | Acetaldehyde | 0.75 | 0.0000 | 14.44 | yes | yes | 9 |
| 8 | 2408-20-0 | Allyl propionate | 0.66 | 0.0000 | 16.09 | yes | yes | 10 |
| 9 | 110-19-0 | Isobutyl acetate | 0.58 | 0.0000 | 19.90 | yes | yes | 11 |
| 10 | 106-36-5 | Propyl propionate | 0.55 | 0.0000 | 18.28 | yes | yes | 20 |
| 11 | 123-86-4 | Butyl acetate | 0.50 | 0.0000 | 11.26 | yes | yes | 13 |
| 12 | 64-19-7 | Acetic acid | 0.48 | 1.0000 | 0.00 | yes | - | 1 |
| 13 | 5878-19-3 | Methoxyacetone | 0.47 | 0.0000 | 8.16 | yes | - | 4 |
| 14 | 123-99-9 | Azelaic acid | 0.44 | 1.0000 | 0.00 | - | - | 1 |
| 15 | 16491-36-4 | cis-3-Hexenyl butyrate | 0.40 | 1.0000 | 0.00 | - | - | 1 |
| 16 | 109-94-4 | Ethyl formate | 0.37 | 0.0000 | 13.29 | yes | - | 9 |
| 17 | 108-62-3 | Metaldehyde | 0.37 | 1.0000 | 0.00 | - | - | 1 |
| 18 | 123-63-7 | Paraldehyde | 0.31 | 0.0000 | 8.80 | - | yes | 5 |
| 19 | 78-98-8 | Pyruvaldehyde | 0.30 | 0.0000 | 14.65 | yes | yes | 13 |
| 20 | 140-88-5 | Ethyl acrylate | 0.29 | 0.0000 | 5.90 | yes | yes | 11 |
| 21 | 19089-92-0 | Hexyl trans-2-butenoate | 0.28 | 1.0000 | 0.00 | - | - | 1 |
| 22 | 7493-71-2 | Allyl tiglate | 0.27 | 0.0000 | 8.56 | yes | yes | 17 |
| 23 | 94133-92-3 | 1-Ethylhexyl tiglate | 0.27 | 0.0000 | 10.86 | - | - | 5 |
| 24 | 2349-13-5 | Heptyl isobutyrate | 0.24 | 1.0000 | 0.00 | - | - | 1 |
| 25 | 540-42-1 | Isobutyl propionate | 0.23 | 0.0000 | 5.63 | yes | yes | 9 |
| 26 | 108-21-4 | Isopropyl acetate | 0.23 | 0.0000 | 4.60 | yes | yes | 8 |
| 27 | 4437-51-8 | 3,4-Hexanedione | 0.23 | 1.0000 | 0.00 | - | - | 1 |
| 28 | 41519-18-0 | Isoamyl tiglate | 0.22 | 0.0000 | 6.34 | - | - | 4 |
| 29 | 7778-87-2 | Propyl heptanoate | 0.22 | 1.0000 | 0.00 | - | - | 1 |
| 30 | 123-92-2 | Isoamyl acetate | 0.22 | 0.0000 | 5.19 | yes | - | 13 |
| 31 | 109-21-7 | Butyl butyrate | 0.21 | 0.0036 | 2.45 | - | - | 7 |
| 32 | 2051-78-7 | Allyl butyrate | 0.20 | 0.0040 | 2.40 | yes | - | 5 |

| | | | | | | | | |
|---|---|---|---|---|---|---|---|---|
| 33 | 105-66-8 | Propyl butyrate | 0.19 | 0.0001 | 3.87 | yes | - | 8 |
| 34 | 3848-24-6 | 2,3-Hexanedione | 0.17 | 0.0046 | 2.34 | - | yes | 5 |
| 35 | 2639-63-6 | Hexyl butyrate | 0.17 | 0.0000 | 5.34 | - | - | 5 |
| 36 | 61692-84-0 | Isobutyl tiglate | 0.17 | 0.0000 | 4.46 | - | - | 5 |
| 37 | 592-84-7 | Butyl formate | 0.17 | 1.0000 | 0.00 | - | - | 9 |
| 38 | 79-09-4 | Propionic acid | 0.16 | 0.0109 | 1.96 | yes | - | 8 |
| 39 | 540-18-1 | amyl butyrate, mixture of isomers | 0.16 | 0.0001 | 4.17 | - | - | 5 |
| 40 | 96-04-8 | 2,3-Heptanedione | 0.16 | 0.0000 | 4.93 | - | yes | 10 |
| 41 | 109-79-5 | 1-Butanethiol | 0.15 | 1.0000 | 0.00 | - | yes | 4 |
| 42 | 52089-55-1 | Ethyl (±)-2-hy-droxycaproate | 0.15 | 0.0046 | 2.34 | - | - | 4 |
| 43 | 71-23-8 | Propanol | 0.15 | 1.0000 | 0.00 | - | - | 3 |
| 44 | 503-74-2 | Isovaleric acid | 0.14 | 0.0291 | 1.54 | - | - | 5 |
| 45 | 637-78-5 | Isopropyl propionate | 0.14 | 1.0000 | 0.00 | yes | - | 2 |
| 46 | 19788-49-9 | Ethyl 2-mercaptopropio-nate | 0.14 | 0.0035 | 2.45 | yes | - | 9 |
| 47 | 6342-56-9 | Pyruvic aldehyde dime-thyl acetal | 0.14 | 0.0016 | 2.78 | - | - | 4 |
| 48 | 107-31-3 | Methyl formate | 0.13 | 0.0002 | 3.80 | yes | - | 9 |
| 49 | 1733-25-1 | Isopropyl tiglate | 0.13 | 1.0000 | 0.00 | yes | - | 1 |
| 50 | 623-37-0 | 3-Hexanol | 0.13 | 1.0000 | 0.00 | yes | - | 2 |
| 51 | 431-03-8 | 2,3-Butanedione | 0.12 | 0.0045 | 2.35 | - | yes | 8 |
| 52 | 64-17-5 | Ethanol | 0.12 | 1.0000 | 0.00 | - | - | 3 |
| 53 | 5910-87-2 | trans, trans-2,4-Nonadi-enal | 0.12 | 1.0000 | 0.00 | - | - | 1 |
| 54 | 624-24-8 | Methyl valerate | 0.11 | 1.0000 | 0.00 | - | - | 11 |
| 55 | 591-12-8 | alpha-Angelicalactone | 0.11 | 0.0015 | 2.83 | - | - | 5 |
| 56 | 140-39-6 | p-Tolyl acetate | 0.10 | 1.0000 | 0.00 | yes | - | 1 |
| 57 | 638-11-9 | Isopropyl butyrate | 0.10 | 0.0074 | 2.13 | yes | - | 5 |
| 58 | 107-92-6 | Butyric acid | 0.09 | 1.0000 | 0.00 | yes | - | 2 |
| 59 | 104-21-2 | Anisyl acetate | 0.09 | 1.0000 | 0.00 | - | - | 1 |
| 60 | 600-14-6 | 2,3-Pentanedione | 0.09 | 1.0000 | 0.00 | yes | yes | 6 |
| 61 | 106-27-4 | Isoamyl butyrate | 0.09 | 0.0200 | 1.70 | - | - | 5 |
| 62 | 79-77-6 | beta-Ionone | 0.09 | 1.0000 | 0.00 | - | - | 2 |
| 63 | 35154-45-1 | cis-3-Hexenyl 3-methyl-butanoate | 0.09 | 1.0000 | 0.00 | - | - | 1 |
| 64 | 591-80-0 | 4-Pentenoic acid | 0.08 | 1.0000 | 0.00 | - | - | 4 |
| 65 | 623-17-6 | Furfuryl acetate | 0.08 | 1.0000 | 0.00 | yes | - | 4 |
| 66 | 21835-01-8 | 3-Ethyl-2-hydroxy-2-cy-clopenten-1-one | 0.08 | 1.0000 | 0.00 | - | - | 4 |
| 67 | 591-68-4 | Butyl valerate | 0.08 | 1.0000 | 0.00 | - | - | 5 |
| 68 | 105-58-8 | Diethyl carbonate | 0.08 | 0.0046 | 2.34 | yes | yes | 9 |
| 69 | 112-31-2 | Decanal | 0.08 | 1.0000 | 0.00 | - | - | 1 |
| 70 | 577-16-2 | 2'-Methylacetophenone | 0.08 | 1.0000 | 0.00 | - | - | 1 |
| 71 | 539-82-2 | Ethyl valerate | 0.08 | 0.0310 | 1.51 | - | yes | 13 |
| 72 | 3268-49-3 | 3-(Methylthio)propio-naldehyde | 0.08 | 1.0000 | 0.00 | - | - | 5 |

| | | | | | | | | |
|---|---|---|---|---|---|---|---|---|
| 73 | 1334-82-3 | Amyl 2-furoate | 0.08 | 1.0000 | 0.00 | - | - | 4 |
| 74 | 2497-18-9 | trans-2-Hexenyl acetate | 0.08 | 0.0006 | 3.21 | - | yes | 9 |
| 75 | 97-53-0 | Eugenol | 0.08 | 1.0000 | 0.00 | - | - | 2 |
| 76 | 97-97-2 | Chloroacetaldehyde dimethyl acetal | 0.07 | 1.0000 | 0.00 | - | - | 4 |
| 77 | 623-19-8 | Furfuryl propionate | 0.07 | 1.0000 | 0.00 | - | - | 4 |
| 78 | 123-75-1 | Pyrrolidine | 0.07 | 1.0000 | 0.00 | - | - | 1 |
| 79 | 4455-13-4 | Ethyl (methylthio)acetate | 0.07 | 1.0000 | 0.00 | - | yes | 5 |
| 80 | 116-53-0 | (±)-2-Methylbutyric acid | 0.07 | 1.0000 | 0.00 | - | - | 4 |
| 81 | 646-07-1 | 4-Methylvaleric acid | 0.07 | 0.0346 | 1.46 | - | - | 4 |
| 82 | 96-48-0 | gamma-Butyrolactone | 0.07 | 1.0000 | 0.00 | yes | - | 9 |
| 83 | 1759-28-0 | 4-Methyl-5-vinylthiazole | 0.07 | 1.0000 | 0.00 | - | - | 1 |
| 84 | 1122-62-9 | 2-Acetylpyridine | 0.06 | 1.0000 | 0.00 | - | - | 1 |
| 85 | 56-86-0 | L-Glutamic acid | 0.06 | 1.0000 | 0.00 | - | - | 1 |
| 86 | 2173-56-0 | Pentyl valerate | 0.05 | 1.0000 | 0.00 | - | - | 5 |
| 87 | 556-82-1 | 3-Methyl-2-buten-1-ol | 0.05 | 1.0000 | 0.00 | - | - | 1 |
| 88 | 142-62-1 | Hexanoic acid | 0.05 | 1.0000 | 0.00 | - | - | 5 |
| 89 | 112-14-1 | Octyl acetate | 0.05 | 1.0000 | 0.00 | - | - | 4 |
| 90 | 589-82-2 | 3-Heptanol | 0.05 | 1.0000 | 0.00 | - | - | 4 |
| 91 | 7764-50-3 | d-Dihydrocarvone | 0.05 | 1.0000 | 0.00 | - | - | 1 |
| 92 | 547-63-7 | Methyl isobutyrate | 0.05 | 1.0000 | 0.00 | yes | - | 6 |
| 93 | 25680-58-4 | 2-Ethyl-3-methoxypyrazine | 0.05 | 1.0000 | 0.00 | - | yes | 9 |
| 94 | 97-64-3 | Ethyl lactate | 0.04 | 0.0339 | 1.47 | yes | - | 7 |
| 95 | 10094-34-5 | alpha,alpha-Dimethylphenethyl butyrate | 0.04 | 1.0000 | 0.00 | - | - | 1 |
| 96 | 3391-87-5 | (+)-Menthone | 0.04 | 1.0000 | 0.00 | - | - | 1 |
| 97 | 106-70-7 | Methyl hexanoate | 0.04 | 1.0000 | 0.00 | - | - | 5 |
| 98 | 7452-79-1 | Ethyl 2-methylbutyrate | 0.04 | 1.0000 | 0.00 | yes | yes | 9 |
| 99 | 108-29-2 | gamma-Valerolactone | 0.04 | 1.0000 | 0.00 | yes | - | 9 |
| 100 | 106-23-0 | (±)-Citronellal | 0.04 | 1.0000 | 0.00 | - | yes | 6 |
| 101 | 111-11-5 | Methyl octanoate | 0.03 | 1.0000 | 0.00 | - | - | 10 |
| 102 | 7785-26-4 | (-)-alpha-Pinene | 0.03 | 1.0000 | 0.00 | - | - | 1 |
| 103 | 103-58-2 | 3-Phenylpropyl isobutyrate | 0.03 | 1.0000 | 0.00 | - | - | 1 |
| 104 | 106-24-1 | Geraniol | 0.03 | 1.0000 | 0.00 | - | - | 4 |
| 105 | 5405-41-4 | Ethyl 3-hydroxybutyrate | 0.03 | 0.0352 | 1.45 | - | - | 7 |
| 106 | 623-42-7 | Methyl butyrate | 0.03 | 1.0000 | 0.00 | yes | yes | 12 |
| 107 | 111-88-6 | 1-Octanethiol | 0.03 | 1.0000 | 0.00 | - | - | 1 |
| 108 | 687-47-8 | (-)-Ethyl L-Lactate | 0.03 | 1.0000 | 0.00 | - | - | 1 |
| 109 | 628-63-7 | Pentyl acetate | 0.03 | 0.0231 | 1.64 | yes | yes | 9 |
| 110 | 2705-87-5 | Allyl cyclohexanepropionate | 0.03 | 1.0000 | 0.00 | - | - | 9 |
| 111 | 1188-02-9 | 2-Methylheptanoic acid | 0.03 | 1.0000 | 0.00 | - | - | 5 |
| 112 | 5454-19-3 | Decyl propionate | 0.03 | 1.0000 | 0.00 | - | - | 2 |
| 113 | 7440-37-1 | Argon (Control) | 0.03 | 1.0000 | 0.00 | - | yes | 44 |

| | | | | | | | | |
|---|---|---|---|---|---|---|---|---|
| 114 | 91-22-5 | Quinoline | 0.03 | 1.0000 | 0.00 | - | - | 1 |
| 115 | 495-40-9 | n-Butyrophenone | 0.03 | 1.0000 | 0.00 | - | - | 1 |
| 116 | 64-04-0 | 2-Phenethylamine | 0.03 | 1.0000 | 0.00 | - | - | 1 |
| 117 | 1797-74-6 | Allyl phenylacetate | 0.03 | 1.0000 | 0.00 | - | - | 1 |
| 118 | 95-92-1 | Diethyl oxalate | 0.03 | 1.0000 | 0.00 | yes | yes | 8 |
| 119 | 695-06-7 | gamma-Hexalactone | 0.02 | 1.0000 | 0.00 | yes | - | 9 |
| 120 | 5146-66-7 | 3,7-Dimethyl-2,6-octadienenitrile | 0.02 | 1.0000 | 0.00 | - | - | 1 |
| 121 | 542-55-2 | Isobutyl formate | 0.02 | 1.0000 | 0.00 | yes | yes | 13 |
| 122 | 123-11-5 | p-Anisaldehyde | 0.02 | 1.0000 | 0.00 | - | - | 1 |
| 123 | 98-01-1 | 2-Furaldehyde | 0.02 | 1.0000 | 0.00 | - | - | 5 |
| 124 | 541-85-5 | 5-Methyl-3-heptanone | 0.02 | 1.0000 | 0.00 | - | - | 1 |
| 125 | 141-05-9 | Diethyl maleate | 0.02 | 1.0000 | 0.00 | - | - | 2 |
| 126 | 104-67-6 | gamma-Undecanolactone | 0.02 | 1.0000 | 0.00 | - | - | 1 |
| 127 | 103-82-2 | Phenylacetic acid | 0.02 | 1.0000 | 0.00 | - | - | 1 |
| 128 | 10031-92-2 | Ethyl 2-nonynoate | 0.01 | 1.0000 | 0.00 | - | - | 4 |
| 129 | 79-31-2 | Isobutyric acid | 0.01 | 1.0000 | 0.00 | - | - | 1 |
| 130 | 89-79-2 | (-)-Isopulegol | 0.01 | 1.0000 | 0.00 | - | - | 1 |
| 131 | 119-36-8 | Methyl salicylate | 0.01 | 1.0000 | 0.00 | - | - | 1 |
| 132 | 105-90-8 | Geranyl propionate | 0.01 | 1.0000 | 0.00 | - | - | 1 |
| 133 | 106-65-0 | Dimethyl succinate | 0.01 | 1.0000 | 0.00 | - | - | 2 |
| 134 | 106-73-0 | Methyl heptanoate | 0.01 | 1.0000 | 0.00 | - | - | 5 |
| 135 | 123-68-2 | Allyl hexanoate | 0.01 | 1.0000 | 0.00 | - | - | 6 |
| 136 | 34047-39-7 | 4-Methylthio-2-butanone | 0.01 | 1.0000 | 0.00 | - | - | 9 |
| 137 | 6728-26-3 | trans-2-Hexenal | 0.00 | 1.0000 | 0.00 | - | - | 1 |
| 138 | 288-47-1 | Thiazole | 0.00 | 1.0000 | 0.00 | - | - | 4 |
| 139 | 540-07-8 | Amyl hexanoate | 0.00 | 1.0000 | 0.00 | - | - | 1 |
| 140 | 60-12-8 | 2-Phenylethyl alcohol | 0.00 | 1.0000 | 0.00 | - | yes | 7 |
| 141 | 142-92-7 | Hexyl acetate | 0.00 | 1.0000 | 0.00 | - | - | 9 |
| 142 | 35466-83-2 | Allyl methyl carbonate | 0.00 | 0.0327 | 1.49 | yes | yes | 9 |
| 143 | 106-32-1 | Ethyl caprylate | 0.00 | 0.0246 | 1.61 | - | - | 10 |
| 144 | 589-38-8 | 3-Hexanone | 0.00 | 1.0000 | 0.00 | - | yes | 5 |
| 145 | 764-48-7 | Ethylene glycol vinyl ether | -0.01 | 1.0000 | 0.00 | - | - | 4 |
| 146 | 93-58-3 | Methyl benzoate | -0.01 | 0.0170 | 1.77 | yes | yes | 31 |
| 147 | 5454-28-4 | Butyl heptanoate | -0.01 | 1.0000 | 0.00 | - | - | 1 |
| 148 | 67883-79-8 | cis-3-Hexenyl tiglate | -0.01 | 1.0000 | 0.00 | - | - | 4 |
| 149 | 40015-15-4 | (Methylthio)acetaldehyde dimethyl acetal | -0.01 | 1.0000 | 0.00 | - | - | 4 |
| 150 | 13679-61-3 | Methyl 2-thiofuroate | -0.01 | 1.0000 | 0.00 | - | - | 4 |
| 151 | 89-82-7 | (R)-(+)-Pulegone | -0.01 | 1.0000 | 0.00 | - | yes | 5 |
| 152 | 106-22-9 | Citronellol | -0.01 | 1.0000 | 0.00 | - | - | 1 |
| 153 | 590-86-3 | 3-Methylbutyraldehyde | -0.01 | 0.0220 | 1.66 | - | yes | 6 |
| 154 | 107-75-5 | 3,7-Dimethyl-7-hydroxyoctanal | -0.02 | 1.0000 | 0.00 | - | - | 1 |

| | | | | | | | | |
|---|---|---|---|---|---|---|---|---|
| 155 | 105-68-0 | Isoamyl propionate | -0.02 | 1.0000 | 0.00 | - | - | 1 |
| 156 | 7492-70-8 | Butyl butyryllactate | -0.02 | 1.0000 | 0.00 | - | - | 1 |
| 157 | 98-86-2 | Acetophenone | -0.02 | 0.0362 | 1.44 | - | - | 7 |
| 158 | 123-51-3 | 3-Methylbutanol | -0.02 | 1.0000 | 0.00 | - | - | 1 |
| 159 | 106-30-9 | Ethyl heptanoate | -0.02 | 1.0000 | 0.00 | - | - | 5 |
| 160 | 629-14-1 | Ethylene glycol diethyl ether | -0.02 | 1.0000 | 0.00 | - | - | 1 |
| 161 | 138-86-3 | Dipentene | -0.02 | 0.0351 | 1.45 | - | - | 3 |
| 162 | 78-70-6 | Linalool | -0.03 | 1.0000 | 0.00 | - | - | 4 |
| 163 | 93-89-0 | Ethyl benzoate | -0.03 | 1.0000 | 0.00 | - | - | 2 |
| 164 | 122-70-3 | 2-Phenylethyl propionate | -0.03 | 1.0000 | 0.00 | - | - | 1 |
| 165 | 15707-24-1 | 2,3-Diethylpyrazine | -0.03 | 1.0000 | 0.00 | - | - | 1 |
| 166 | 105-87-3 | Geranyl acetate | -0.03 | 1.0000 | 0.00 | - | - | 1 |
| 167 | 105-54-4 | Ethyl butyrate | -0.03 | 1.0000 | 0.00 | yes | - | 8 |
| 168 | 6622-76-0 | Methyl tiglate | -0.03 | 0.0060 | 2.22 | - | yes | 12 |
| 169 | 15707-23-0 | 2-Ethyl-3-methylpyrazine | -0.04 | 1.0000 | 0.00 | - | - | 1 |
| 170 | 7540-53-6 | Citronellyl valerate | -0.04 | 0.0070 | 2.15 | - | - | 5 |
| 171 | 300-57-2 | Allyl benzene | -0.04 | 1.0000 | 0.00 | - | - | 3 |
| 172 | 137-00-8 | 4-Methyl-5-thiazoleethanol | -0.04 | 1.0000 | 0.00 | - | - | 1 |
| 173 | 623-70-1 | Ethyl crotonate | -0.04 | 1.0000 | 0.00 | yes | - | 5 |
| 174 | 53448-07-0 | trans-2-Undecenal | -0.04 | 1.0000 | 0.00 | - | - | 1 |
| 175 | 67-56-1 | Methanol | -0.05 | 1.0000 | 0.00 | - | - | 3 |
| 176 | 624-48-6 | Dimethyl maleate | -0.05 | 1.0000 | 0.00 | yes | - | 1 |
| 177 | 108-98-5 | Thiophenol | -0.05 | 1.0000 | 0.00 | - | - | 1 |
| 178 | 123-66-0 | Ethyl hexanoate | -0.05 | 0.0045 | 2.35 | - | yes | 14 |
| 179 | 1009-14-9 | Valerophenone | -0.05 | 1.0000 | 0.00 | - | - | 1 |
| 180 | 123-32-0 | 2,5-Dimethylpyrazine | -0.05 | 1.0000 | 0.00 | - | - | 2 |
| 181 | 589-98-0 | 3-Octanol | -0.05 | 1.0000 | 0.00 | - | - | 1 |
| 182 | 97-99-4 | Tetrahydrofurfuryl alcohol | -0.05 | 0.0142 | 1.85 | - | - | 5 |
| 183 | 470-67-7 | Cineole | -0.06 | 0.0286 | 1.54 | - | yes | 7 |
| 184 | 100-52-7 | Benzaldehyde | -0.06 | 0.0002 | 3.66 | - | - | 6 |
| 185 | 107-87-9 | 2-Pentanone | -0.06 | 1.0000 | 0.00 | - | yes | 12 |
| 186 | 1128-08-1 | Dihydrojasmone | -0.06 | 1.0000 | 0.00 | - | - | 1 |
| 187 | 78-93-3 | 2-Butanone | -0.07 | 0.0234 | 1.63 | yes | - | 3 |
| 188 | 925-78-0 | 3-Nonanone | -0.09 | 0.0184 | 1.73 | - | - | 2 |
| 189 | 543-49-7 | 2-Heptanol | -0.09 | 0.0049 | 2.31 | - | - | 4 |
| 190 | 2396-83-0 | Ethyl 3-hexenoate | -0.09 | 1.0000 | 0.00 | - | - | 1 |
| 191 | 51729-83-0 | Methyl isopropyl carbonate | -0.09 | 0.0000 | 5.67 | - | - | 5 |
| 192 | 562-74-3 | Terpinen-4-ol | -0.09 | 1.0000 | 0.00 | - | - | 1 |
| 193 | 6976-93-8 | Ethylene glycol methyl ether methacrylate | -0.10 | 0.0001 | 4.01 | - | - | 4 |
| 194 | 591-78-6 | 2-Hexanone | -0.10 | 0.0002 | 3.64 | - | - | 6 |
| 195 | 590-01-2 | Butyl propionate | -0.11 | 0.0021 | 2.68 | yes | yes | 5 |

| | | | | | | | | |
|---|---|---|---|---|---|---|---|---|
| 196 | 629-19-6 | Propyl disulfide | -0.11 | 0.0067 | 2.17 | - | - | 4 |
| 197 | 108-48-5 | 2,6-Lutidine | -0.11 | 0.0002 | 3.74 | - | - | 4 |
| 198 | 20487-40-5 | tert-Butyl propionate | -0.11 | 0.0076 | 2.12 | - | - | 4 |
| 199 | 112-30-1 | 1-Decanol | -0.11 | 0.0002 | 3.72 | - | - | 4 |
| 200 | 110-93-0 | 6-Methyl-5-hepten-2-one | -0.13 | 1.0000 | 0.00 | - | - | 1 |
| 201 | 5837-78-5 | Ethyl tiglate | -0.13 | 0.0006 | 3.19 | yes | - | 13 |
| 202 | 110-62-3 | Valeraldehyde | -0.13 | 1.0000 | 0.00 | - | - | 1 |
| 203 | 623-36-9 | 2-Methyl-2-pentenal | -0.13 | 0.0000 | 5.10 | - | yes | 24 |
| 204 | 100-66-3 | Anisole | -0.13 | 0.0000 | 4.66 | - | - | 18 |
| 205 | 111-13-7 | 2-Octanone | -0.14 | 1.0000 | 0.00 | - | - | 1 |
| 206 | 110-43-0 | 2-Heptanone | -0.15 | 0.0000 | 5.14 | - | - | 5 |
| 207 | 124-13-0 | Octanal | -0.15 | 0.0000 | 5.71 | - | - | 3 |
| 208 | 25152-84-5 | trans, trans-2,4-Decadienal | -0.15 | 0.0063 | 2.20 | - | - | 4 |
| 209 | 68480-28-4 | 3-Methylbut-2-enyl formate | -0.17 | 0.0000 | 6.78 | - | - | 4 |
| 210 | 19700-21-1 | Geosmin | -0.20 | 1.0000 | 0.00 | - | - | 1 |
| 211 | 111-71-7 | Heptanal | -0.28 | 1.0000 | 0.00 | - | - | 1 |
| 212 | 121-45-9 | Trimethyl phosphite | -0.32 | 0.0000 | 10.99 | - | - | 4 |
| 213 | 66-25-1 | Hexanal | -0.33 | 1.0000 | 0.00 | - | - | 1 |

**Supplementary Table ST2:** Abbreviations of Descriptor Blocks as used in the manuscript.

| **Abbreviation** | **Descriptors** |
|---|---|
| #FGROUP | FUNCTIONAL GROUP COUNTS |
| 2DAUTO | TWOD AUTOCORRELATIONS |
| 3DMORSE | THREEDMORSE |
| A_FRAG | ATOMCENTRED FRAGMENTS |
| BCUT | BURDEN EIGENVALUES DESCRIPTORS |
| CONST | CONSTITUTIONAL |
| C_IND | CONNECTIVITY INDICES |
| EA_IND | EDGE ADJACENCY INDICES |
| EV_IND | EIGENVALUE INDICES |
| GEO | GEOMETRICAL |
| GETAWAY | GETAWAY |
| MOL_PROP | MOLECULAR PROPERTIES |
| RAND | RANDIC MOLECULAR PROFILES |
| RDF | RDF |
| TOPO | TOPOLOGICAL |
| WHIM | WHIM |
| WPATH | WALK PATH COUNTS |
